# *GBA1* deficiency differentially affects endolysosomal trafficking in neurons versus astrocytes

**DOI:** 10.64898/2026.01.12.697563

**Authors:** Anna J. Park, Sarah L. Fish, Colby L. Samstag, Minsuh Kim, Jeremy Weiss, Selina Yu, Malia L. Callier, Ella H. Chiu, Joshua Weiss, Christiane S. Hampe, Raja E. Estes, Bernice Lin, Arnav Khera, Dora Yearout, Leo J. Pallanck, Cyrus P. Zabetian, Marie Y. Davis

**Author notes:** Corresponding author: Marie Y. Davis, MD, PhD, VA Puget Sound Healthcare System 1660 S. Columbian Way, Seattle, WA 98108. These authors contributed equally (Anna J. Park, Sarah L. Fish). Albert Einstein College of Medicine, Bronx, NY. University of Montana, Missoula, MT. The Johns Hopkins University School of Medicine, Baltimore, MD.

## Abstract

Mutations in the gene *glucosidase, beta acid 1 (GBA1)* are the strongest genetic risk factor for Parkinson’s disease (PD) and are associated with faster disease progression. *GBA1* is expressed in all cell types of the central nervous system, with some evidence supporting higher expression in glial cells than neurons. To elucidate possible specific functions in neurons versus glia, we differentiated human induced pluripotent stem cells (iPSCs) generated from an individual with PD heterozygous for the *GBA1* pathogenic null variant IVS2+1 (*GBA1^IVS/+^*), homozygous *GBA1* IVS2+1 isogenic to G*BA1^IVS/+^* (*GBA1^IVS/IVS^*) and a healthy unaffected age-and sex-matched individual (*GBA1^+/+^*).

*GBA1* expression was reduced in *GBA1^IVS/+^* and *GBA1^IVS/IVS^* neurons and astrocytes.

Endolysosomal trafficking was significantly altered in G*BA1-*deficient neurons with enlarged early and recycling endosome and lysosome compartments in neurons but not in astrocytes. High molecular weight oligomerization of α-synuclein and phosphorylated Ser129 α-synuclein were present in *GBA1^IVS/+^* and *GBA1^IVS/IVS^* neurons but not in *GBA1^+/+^* neurons, or in *GBA1-*deficient or *GBA1^+/+^* astrocytes. Transcriptomic analysis of *GBA1*-deficient neurons and astrocytes revealed cell-type specific profiles. *GBA1* deficiency in neurons downregulated immune response and upregulated cholesterol synthesis pathways, while *GBA1* deficiency in astrocytes downregulated genes associated with translation and upregulated genes involved in extracellular matrix biogenesis. Transcriptomic analysis also suggests that *GBA1* deficiency induces neurotoxic reactivity in astrocytes. Together, these findings indicate that *GBA1* deficiency has cell type-specific effects, with increased neuronal vulnerability to endolysosomal trafficking leading to α-synucleinopathy while *GBA1* deficiency in astrocytes leads to increased neurotoxic reactivity independent of endolysosomal trafficking and α-synucleinopathy.

**Highlights:**

- iPSC-derived neurons and astrocytes modeled *GBA1* deficiency
- Endolysosomal trafficking defects occurred only in *GBA1*-deficient neurons
- α-synuclein oligomers accumulated in *GBA1*-deficient neurons, not astrocytes
- Astrocyte *GBA1* loss drove neurotoxic reactive gene signatures
- *GBA1* deficiency causes cell-type specific pathology

## Introduction

α-synucleinopathies, encompassing Parkinson’s disease (PD), dementia with Lewy bodies (DLB), and multiple system atrophy (MSA), are together the second most common group of neurodegenerative disorders following Alzheimer’s disease (AD). Like AD and other neurodegenerative disorders, α-synucleinopathies are characterized by the neuropathological hallmark of intracellular proteinaceous aggregates, specifically α-synuclein oligomers. These aggregates localize intraneuronally in PD and DLB, and are intraglial in MSA. Impairment in multiple cellular processes such as autophagy and mitophagy have been implicated in the process leading to accumulation of aberrant α-synuclein^1,2^. However, the precise mechanisms by which abnormal α-synuclein processing contribute to neurodegeneration remain unclear.

Heterozygous variants in the *glucosylceramidase beta 1 (GBA1)* gene, encoding the lysosomal lipid metabolism enzyme glucocerebrosidase, are the strongest genetic risk factor for PD and DLB, with odds ratios between 3 and 20^3–6^. *GBA1* variants are not uncommon as 5-15% of PD patients are found to carry a *GBA1* variant^6–8^. *GBA1* variants are not only associated with increased risk but also accelerated clinical progression of both motor and cognitive symptoms in PD^9–11^.

Our prior work in a *Drosophila* model of *GBA1* deficiency demonstrated altered extracellular vesicle biogenesis resulting in the accelerated spread of protein aggregation, implicating the endolysosomal trafficking system^12–14^. Endolysosomal trafficking is a complex and highly regulated intracellular vesicular system that encompasses internalization of extracellular cargo targeted to specific organelles, recycling and degradation of intracellular proteins and cargo, a secretion of cargo into the extracellular space, either packaged within membrane bound extracellular vesicles or directly into the extracellular matrix^15^. Impaired endolysosomal trafficking is implicated in the pathogenesis of neurodegenerative diseases, including AD and PD^1,12–14,16,17^ and is largely unexplored in how the processing of pathogenic proteins in different cell types influences neurodegeneration. Interestingly, the efficiency of endolysosomal trafficking is tailored to specific cell types, where cells with phagocytic activity such as microglia and astrocytes have been found to have more efficient endocytic trafficking of cargo, including α-synuclein compared to neurons^2,18^.

Although astrocytes have lower endogenous expression of α-synuclein^19^, α-synucleinopathy is found in both neurons and astrocytes in PD and DLB, and predominantly in glia in MSA^20^. *In vitro* studies have demonstrated that astrocytes can internalize a-synuclein oligomers more rapidly than neurons^2,21,22^, and network analysis of proteomic data suggests that astrocytic endolysosomal processing of α-synuclein differs from neuronal processing^23^. These studies suggest that endolysosomal trafficking can differ in capacity in specific cell types, which could have implications on cell type-specific susceptibility to neurodegeneration.

To investigate whether *GBA1* deficiency has a differential effect on endolysosomal trafficking in neurons and glia, we generated both dopaminergic midbrain neurons and astrocytes from control and *GBA1-*deficient human iPSCs carrying a null mutation.

Interestingly, we found that *GBA1* deficiency had a significant impact on early endosome, recycling endosome, and lysosome morphology in neurons but not astrocytes. The protein expression of early endosome antigen 1 (EEA1*)-* a marker for early endosomes and Rab11*-* a small GTPase protein involved in vesicular trafficking and recycling endosomes, was also increased in *GBA1-*deficient neurons but not astrocytes. The expression of lysosome-associated membrane protein 1 (LAMP1)-a protein mainly found on lysosome and late endosomes, was increased in *GBA1*-deficient neurons and astrocytes with complete knockout of *GBA1*. Pathologic phosphorylated and oligomeric α-synuclein was only detected in *GBA1-*deficient neurons, not astrocytes. These results indicate that *GBA1-*deficiency perturbs endolysosomal trafficking and α-synuclein processing in a cell-specific manner.

Transcriptomic profiling of *GBA1*-deficient neurons and astrocytes surprisingly revealed minimal overlap of differentially expressed genes in astrocytes and neurons, indicating more divergent cell-type specific alterations than expected. Expression of genes involved in cholesterol metabolism and immune response were specifically altered in *GBA1*-deficient neurons, while genes involved in extracellular matrix and translation were specifically dysregulated in astrocytes. Together, our findings indicate that *GBA1* deficiency drives specific disease-relevant pathogenic processes in neurons and astrocytes, linking impaired endolysosomal trafficking to α-synucleinopathy in neurons, and promoting reactive transcriptional changes in astrocytes independent of α-synucleinopathy in astrocytes. Elucidating how *GBA1* deficiency impacts cell-specific functions of neurons and glia is critical for better understanding the complex pathogenesis of α-synucleinopathies and developing novel therapeutic cell-type specific strategies to mitigate neurodegeneration.

## Results

We generated human iPSCs from fibroblasts from an individual with PD carrying the *GBA1* c.115+1 G>A, also known as IVS2+1 G>A variant (*GBA1^IVS/+^*). This intronic *GBA1* variant is associated with severe neuronopathic Gaucher’s disease and interrupts the canonical splice site following exon 2, leading to nonsense mediated decay and is therefore considered a null allele^24^. An isogenic *GBA1^+/+^ Rev* control was generated by CRISPR/Cas9 genome editing to revert the *GBA1* IVS2+1 G>A variant back to wildtype, and in the genome editing process, an iPSC clone homozygous for *GBA1* IVS2+1 G>A was also generated (*GBA1^IVS/IVS^*) (S Fig 1). A matched control iPSC clone that remained unedited after CRISPR/Cas9 genome editing, maintaining heterozygosity for *GBA1* IVS2+1 G>A, was also isolated (*GBA1^IVS/+^* A4) (Fig 1A). All isolated lines generated by CRISPR/Cas9 editing were confirmed to not have alterations to the pseudogene *GBAP1* (S. Fig 1). Additional control iPSC lines were generated from peripheral blood monocytes from a healthy age-and sex-matched individual (*GBA1^+/+^*) unrelated to the individual carrying the *GBA1* IVS2+1 G>A variant.

**Figure 1.**
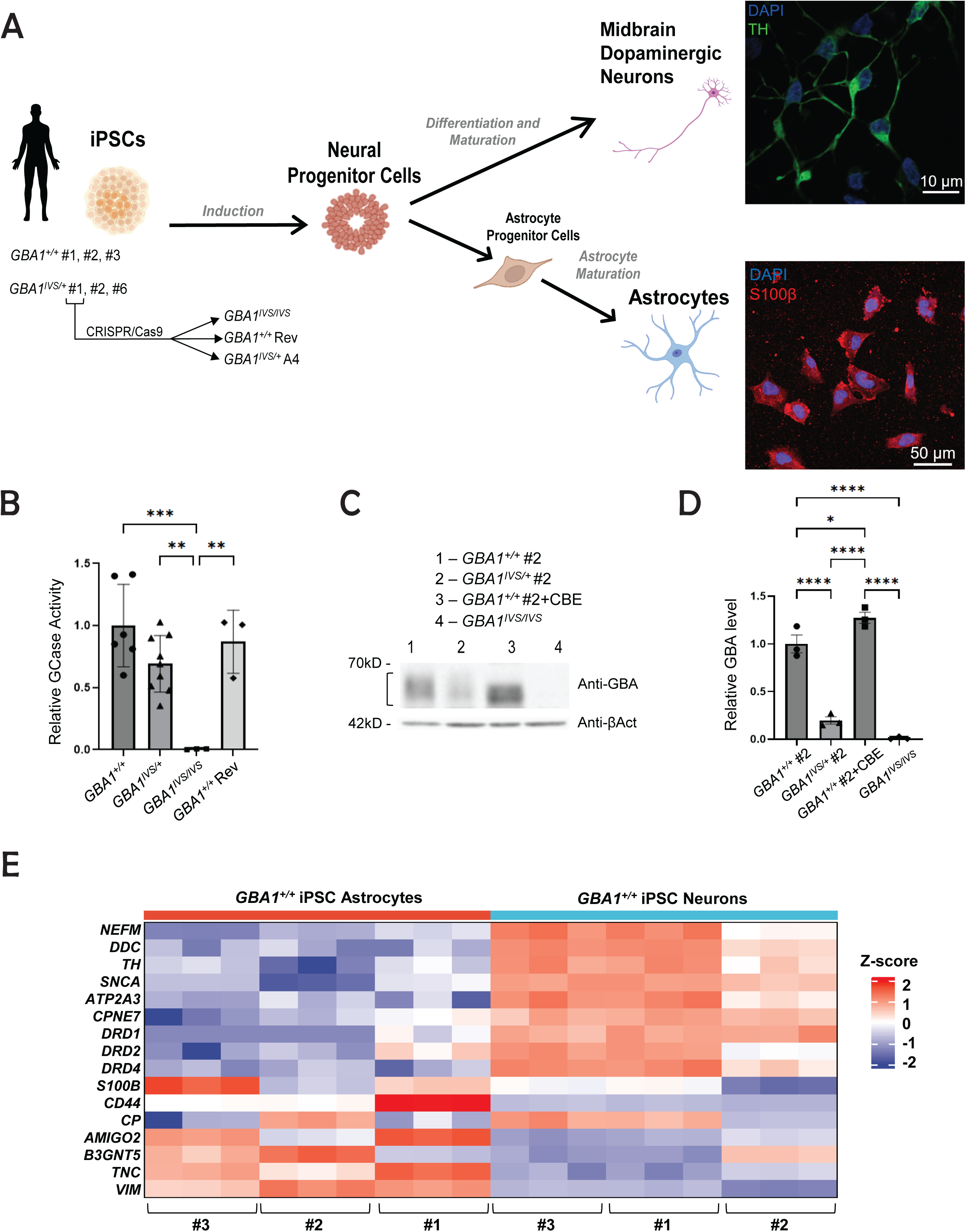
Differentiation of *GBA1-deficient* iPSCs into midbrain dopaminergic neurons and astrocytes. **(A)** Diagram of iPSC cell line generation and differentiation protocol into midbrain dopaminergic neurons and astrocytes, with **r**epresentative images of immunocytochemistry of *GBA1^+/+^* iPSC-neurons stained with anti-TH (green) and DAPI and *GBA1^+/+^* iPSC-astrocytes stained with anti-S100β (red) and DAPI **(B)** Quantification of glucocerebrosidase (GCase) enzyme activity by relative fluorescence of 4-methylumbelliferyl-β-D-glucopyranoside (4-MUG) in indicated genotypes. GCase activity of all cell lines treated with CBE was subtracted to eliminate background activity. (one-way ANOVA F (4, 19) =9.092, p=0.0003). **(C)** Representative Western blot with anti-GBA1 protein in *GBA1^+/+^*, *GBA1^IVS/+^*, *GBA1^+/+^* + conduritol B epoxide (CBE), and *GBA1^IVS/IVS^*iPSC-neurons. **(D)** Quantification of GBA1 protein in neurons from 3 independent replicates in indicated genotypes, normalized to Actin in control (one-way ANOVA F (3,8) =108.6, p<0.0001). **(E)** Heatmap of differentially expressed midbrain dopaminergic neuron-and astrocyte-specific genes in *GBA1^+/+^* neurons versus *GBA1^+/+^*astrocytes sorted by z-score. Statistical significance determined by one-way ANOVA (ns = p ≥ 0.05, * = p < 0.05, ** = p < 0.01, *** = p < 0.001, **** = p < 0.0001). Data are presented as mean ± SEM.

**Figure 2.**
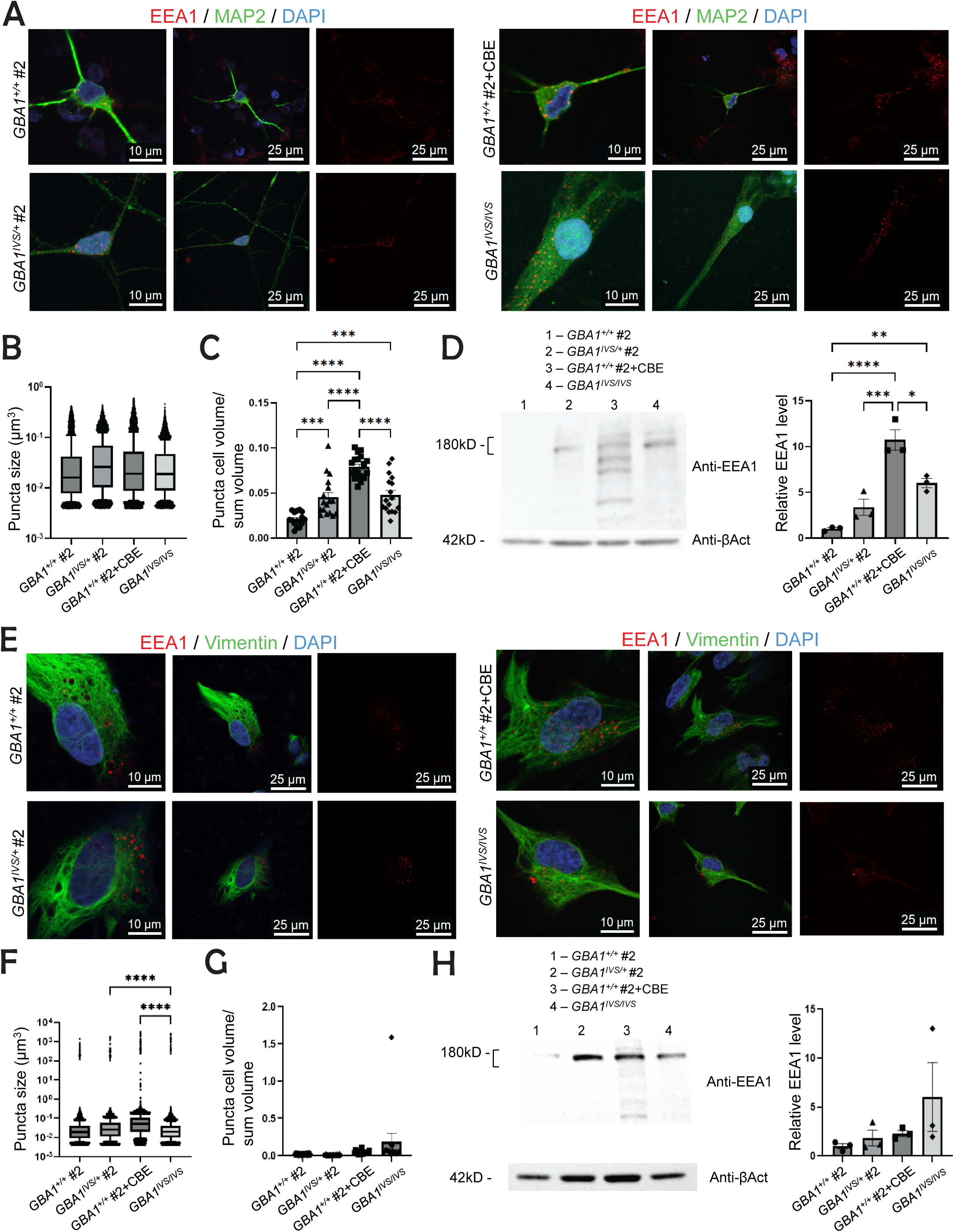
Early endosomes are altered in *GBA1*-deficient neurons, but not in *GBA1*-deficient astrocytes. **(A)** Representative images of immunocytochemistry of anti-EEA1 (red), anti-MAP2 (green) and DAPI in *GBA1^+/+^*, *GBA1^IVS/+^*, *GBA1^+/+^* + conduritol B epoxide (CBE), and *GBA1^IVS/IVS^* neurons. **(B)** Quantification of anti-EEA1 puncta volume per neuron across indicated genotypes (one-way ANOVA: F(3, 29925) = 104.0, p<0.0001). One-way ANOVA did not detect statistically significant differences between groups. Cohen’s *d* effect sizes were calculated, with only medium and large effects displayed. **(C)** Ratio of anti-EEA1 puncta volume to total anti-MAP2 neuronal cell volume in indicated genotypes (F(3, 64) = 34.35, p<0.0001). **(D)** Western blot of anti-EEA1 protein levels in neurons in indicated genotypes, with quantification normalized to Actin in control. **(E)** Representative immunocytochemistry images of anti-EEA1 (red), anti-vimentin (green) and DAPI in *GBA1^+/+^*, *GBA1^IVS/+^*, *GBA1^+/+^* + conduritol B epoxide (CBE), and *GBA1^IVS/IVS^*astrocytes. **(F)** Quantification of anti-EEA1 puncta volume per astrocyte in indicated genotypes (F(3, 35549) = 13.42, p<0.0001). **(G)** Ratio of anti-EEA1 puncta volume to total anti-Vimentin astrocyte cell volume in indicated genotypes (F(3, 65) = 2.761, p=0.0491). **(H)** Western blot of anti-EEA1 protein levels in astrocytes in indicated genotypes, with quantification normalized to Actin in control. Statistical significance was determined by one-way ANOVA (ns = p ≥ 0.05, * = p < 0.05, ** = p < 0.01, *** = p < 0.001, **** = p < 0.0001). For quantification of immunocytochemistry, statistical significance was determined by p < 0.05 and Cohen’s *d* of medium and large effect. Data are presented as mean ± SEM.

Three *GBA1^IVS^*^/+^ clones (#1,2,6), three *GBA1^+/+^* clones (#1,2,3), a *GBA1^+/+^ Rev* isogenic control clone, *GBA1^IVS^*^/+^ A4 clone, and a homozygous null *GBA1^IVS/IVS^* iPSC clone were confirmed to not have chromosomal abnormalities by karyotype differentiated into neural progenitor cells (NPCs) prior to differentiation into midbrain dopaminergic neurons and astrocytes (S. Fig 2). Neuronal differentiation of NPCs was confirmed by anti-microtubule-associated protein 2 (MAP2) expression, and differentiation into dopaminergic neurons was confirmed by anti-tyrosine hydroxylase (TH) staining (Fig 1A). Astrocyte identity was confirmed by anti-S100β (Fig 1A). To confirm reduction in glucocerebrosidase (GCase) enzyme activity in *GBA1^IVS^*^/+^ and *GBA1^IVS/IVS^* NPCs, we measured GCase activity using the fluorescent synthetic glucocerebrosidase substrate 4-methylumbelliferyl-beta-D-glucopyranoside. *GBA1^+/+^* treated with conduritol B epoxide (CBE), an irreversible inhibitor of glucocerebrosidase, was used as a positive control. While *GBA1^IVS/+^* NPCs trended to reduced glucocerebrosidase (GCase) enzyme activity compared to *GBA1^+/+^* clones, GCase enzyme activity was nearly undetectable in *GBA1^IVS/IVS^* NPCs (Fig 1B). GBA1 protein levels were significantly reduced in *GBA1^IVS/+^* neurons and nearly undetectable in *GBA1^IVS/IVS^* neurons (Fig 1C-D).

To further validate that we successfully differentiated iPSCs into midbrain dopaminergic neurons and astrocytes we performed RNA-seq of neurons and astrocytes differentiated from unaffected healthy *GBA1^+/+^* clones. Transcriptomic profiles confirmed significantly increased expression of neuronal-specific genes in control neurons, and astrocyte-specific genes in control astrocytes (Fig 1E). Consistent with known reduced expression of *SNCA* in glia compared to neurons, *SNCA* expression was downregulated in *GBA1^+/+^* astrocytes compared to *GBA1^+/+^* neurons by 73%.

Upregulations of *TH, DDC, DRD1* and *DRD4* in neurons compared to astrocytes is consistent with successful differentiation into midbrain dopaminergic neurons (Fig 1E).

We expected that the transcriptomic profile of cells differentiated from the *GBA1^+/+^ Rev* isogenic control line would be largely concordant the three control *GBA1^+/+^* clonal lines, however we found that over 6,000 genes were differentially expressed between *GBA1^+/+^* and *GBA1^+/+^ Rev* neurons (S Fig 3A). Furthermore, principal component analysis (PCA) (S Fig 3B) revealed significant separation between *GBA1^+/+^* and *GBA1^+/+^ Rev* neurons, with 78.43% total variance in the first principal component and 20.65% total variance in the second principal component. 2,774 genes were differentially regulated between control *GBA1^+/+^* clones and isogenic *GBA1^+/+^ Rev* astrocytes (S Fig 4A). PCA comparing astrocytes from 3 *GBA1^+/+^* clones and isogenic *GBA1^+/+^ Rev* astrocytes revealed significant separation with 53% total variance in the first component and 29.9% total variance in the second component (S Fig 4B). This significant difference in transcriptomic profile between our isogenic *GBA1^+/+^ Rev* and the unrelated age-and sex-matched *GBA1^+/+^* was surprising and led us to suspect that the transcriptomic alterations in the *GBA1^+/+^ Rev* isogenic iPSC control line may be due to the 10 additional passages while undergoing CRISPR/Cas9 editing. To test this hypothesis, we examined the transcriptomic profiles of astrocytes and neurons differentiated from *GBA1^IVS^*^/+^ A4 iPSCs which underwent the CRISPR/Cas9 genome editing process in parallel to the cells that resulted in the isogenic control *GBA1^+/+^ Rev* but remained unedited and therefore should be equivalent to *GBA1^IVS^*^/+^ cells. However, *GBA1^IVS^*^/+^ A4 neurons also differed significantly in transcriptomic profile compared to *GBA1^IVS^*^/+^ neurons, with 2,319 differentially expressed genes, and greater than 50% total variance in the first component of principal component analysis (S Fig 5A). The transcriptomic profiles of *GBA1^IVS^*^/+^ A4 astrocytes compared to *GBA1^IVS^*^/+^ astrocytes were less divergent, with only 585 genes differentially expressed (S Fig 5B). However, because of the concern that the additional passages and CRISPR/Cas9 genome editing had additional adverse effects we used iPSCs generated from an unrelated age-and sex-matched unaffected control and an additional negative control of healthy age-and sex-matched *GBA1^+/+^* cells treated with 100mM Conduritol B epoxide (CBE), an irreversible inhibitor of glucocerebrosidase, in our subsequent analyses.

**Figure 3.**
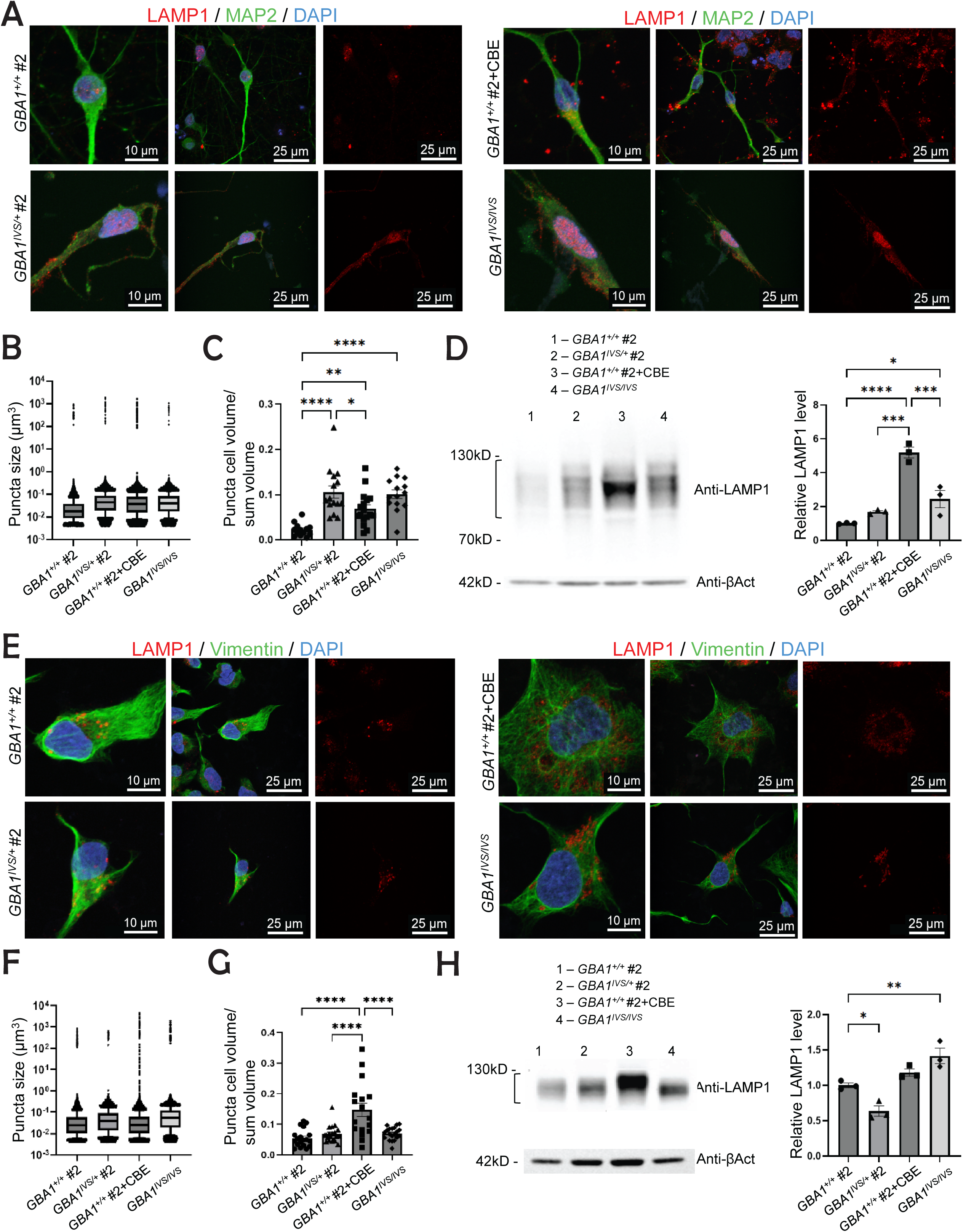
Lysosomes are significantly altered in *GBA1*-deficient neurons, but not in *GBA1*-deficient astrocytes. **(A)** Representative images of immunocytochemistry of anti-LAMP1 (red), anti-MAP2 (green) and DAPI in *GBA1^+/+^*, *GBA1^IVS/+^*, *GBA1^+/+^* + conduritol B epoxide (CBE), and *GBA1^IVS/IVS^* neurons. **(B)** Quantification of anti-LAMP1 puncta volume per neuron across genotypes (one-way ANOVA: F(3, 68096) = 4.555, p=0.0034). **(C)** Ratio of anti-LAMP1 puncta volume to total anti-MAP2 neuronal cell volume across genotypes (F(3, 55) = 13.95, p<0.0001). **(D)** Western blot analysis of LAMP1 protein levels in neurons in indicated genotypes, with quantification normalized to Actin in control. **(E)** Representative images of immunocytochemistry of anti-LAMP1 (red), anti-Vimentin (green) and DAPI in *GBA1^+/+^*, *GBA1^IVS/+^*, *GBA1^+/+^* + conduritol B epoxide (CBE), and *GBA1^IVS/IVS^* astrocytes. **(F)** Quantification of anti-LAMP1 puncta volume per astrocyte across genotypes (F(3, 61687) = 2.274, p=0.0778). **(G)** Ratio of anti-LAMP1 puncta volume to total anti-Vimentin astrocyte cell volume in indicated genotypes (F(3, 72) = 13.48, p<0.0001). **(H)** Western blot analysis of LAMP1 protein level in astrocytes in indicated genotypes, with quantification normalized to Actin in control. Statistical significance was determined by one-way ANOVA (ns = p ≥ 0.05, * = p < 0.05, ** = p < 0.01, *** = p < 0.001, **** = p < 0.0001). For quantification of immunocytochemistry, statistical significance was determined by p < 0.05 and Cohen’s *d* of medium and large effect. Data are presented as mean ± SEM.

**Figure 4.**
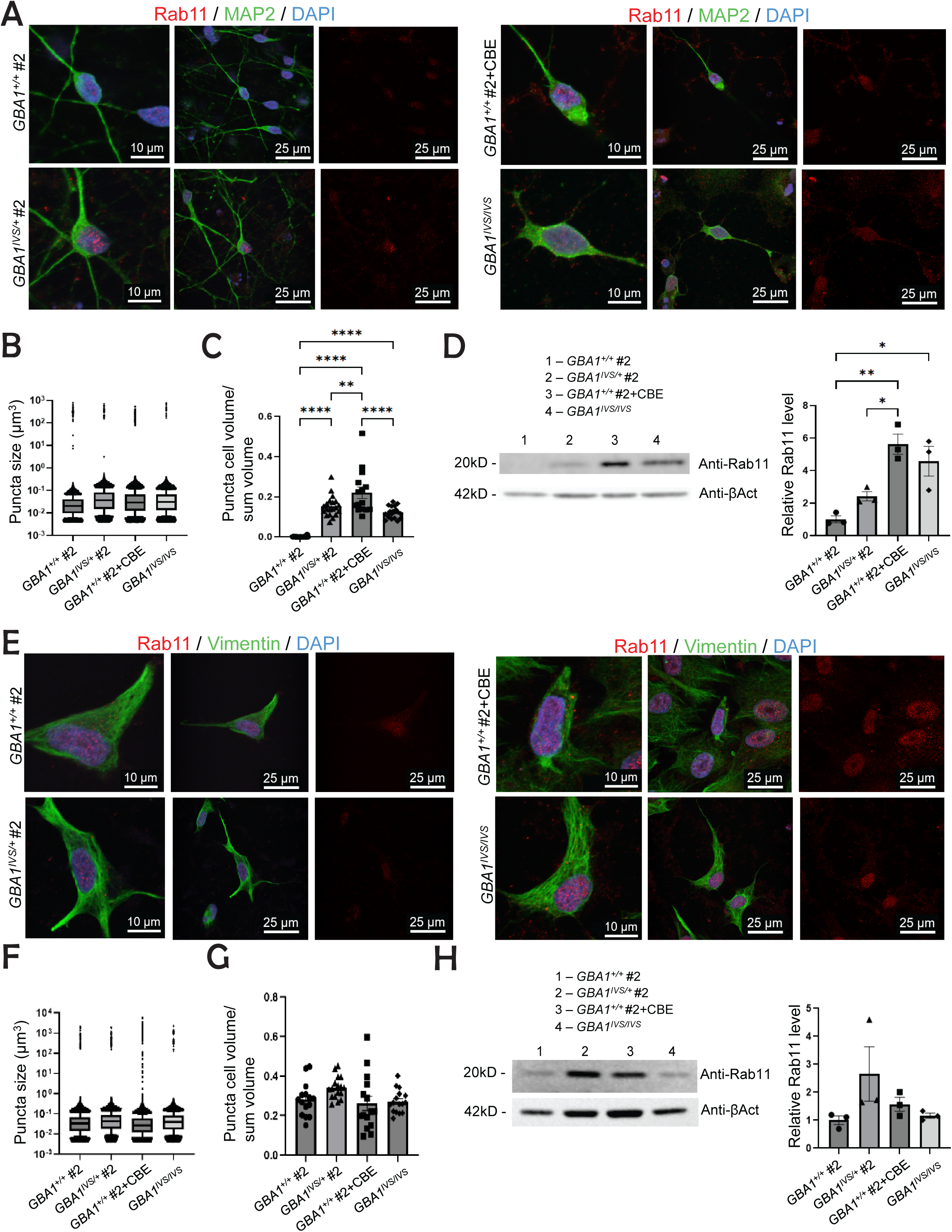
Recycling endosomes are significantly altered in *GBA1-*deficient neurons, but not in *GBA1-*deficient astrocytes. **(A)** Representative images of immunocytochemistry of anti-Rab11 (red), anti-MAP2 (green), and DAPI in *GBA1^+/+^*, *GBA1^IVS/+^*, *GBA1^+/+^* + conduritol B epoxide (CBE), and *GBA1^IVS/IVS^* neurons. **(B)** Quantification of Rab11 puncta volume per neuron in indicated genotypes (one-way ANOVA: F(3, 59810) = 22.77, p<0.0001). **(C)** Ratio of Rab11 puncta volume to total anti-MAP2 neuronal cell volume in indicated genotypes (F(3, 71) = 45.67, p<0.0001). **(D)** Western blot analysis of Rab11 protein levels in neurons across genotypes, with quantification normalized to Actin in control. **(E)** Representative images of immunocytochemistry of anti-Rab11 (red), anti-Vimentin (green) and DAPI in *GBA1^+/+^*, *GBA1^IVS/+^*, *GBA1^+/+^* + conduritol B epoxide (CBE), and *GBA1^IVS/IVS^*astrocytes. **(F)** Quantification of Rab11 puncta volume per astrocyte in indicated genotypes (F(3, 233665) = 0.4953, p=0.6855). **(G)** Ratio of anti-Rab11 puncta volume to total anti-Vimentin astrocyte cell volume in indicated genotypes (F(3, 59) = 2.529, p=0.0659). **(H)** Western blot analysis of Rab11 protein levels in astrocytes across genotypes, with quantification normalized to Actin in control. Statistical significance was determined by one-way ANOVA (ns = p ≥ 0.05, * = p < 0.05, ** = p < 0.01, *** = p < 0.001, **** = p < 0.0001). For quantification of immunocytochemistry, statistical significance was determined by p < 0.05 and Cohen’s *d* of medium and large effect. Data are presented as mean ± SEM.

**Figure 5.**
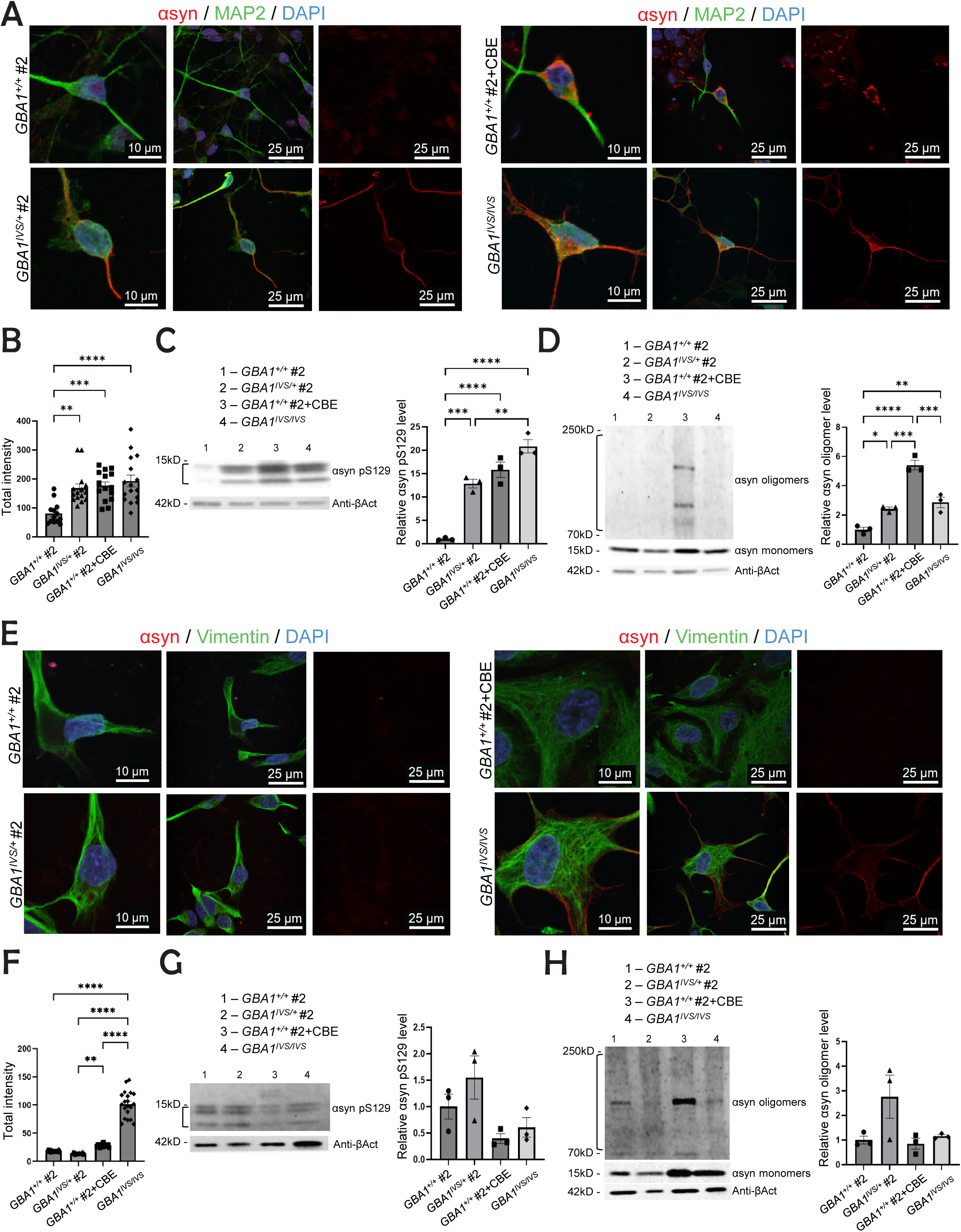
α-syn and pSer129 α-syn levels are significantly altered in *GBA1*-deficient neurons, but not in *GBA1*-deficient astrocytes. **(A)** Representative images of immunocytochemistry of anti-α-syn (red), anti-MAP2 (green) and DAPI in *GBA1^+/+^*, *GBA1^IVS/+^*, *GBA1^+/+^* + conduritol B epoxide (CBE), and *GBA1^IVS/IVS^* neurons. **(B)** Quantification of immunocytochemistry of anti-α-syn total intensity per anti-MAP2 neuron in indicated genotypes (one-way ANOVA: F(3, 55) = 9.560, p<0.0001). **(C)** Western blot analysis of anti-pSer129 α-syn levels in neurons in indicated genotypes, with quantification normalized to Actin in control. **(D)** Western blot analysis of α-syn high molecular weight oligomer (70-250 kD) and α-syn monomer (15 kD) levels in neurons in indicated genotypes, with quantification normalized to Actin in control. **(E)** Representative images of immunocytochemistry of anti-α-syn in *GBA1^+/+^*, *GBA1^IVS/+^*, *GBA1^+/+^* + conduritol B epoxide (CBE), and *GBA1^IVS/IVS^*astrocytes. **(F)** Quantification of immunocytochemistry of anti-α-syn total intensity per anti-Vimentin astrocyte volume in indicated genotypes (F(3, 70) = 220.8, p<0.0001). **(G)** Western blot analysis of pSer129 α-syn levels in astrocytes in indicated genotypes, with quantification normalized to Actin in control. **(H)** Western blot analysis of α-syn oligomer (70-250 kD) and α-syn monomer (15 kD) levels in astrocytes across genotypes, with quantification normalized to Actin in control. Statistical significance was determined by one-way ANOVA (ns = p ≥ 0.05, * = p < 0.05, ** = p < 0.01, *** = p < 0.001, **** = p < 0.0001). For quantification of immunocytochemistry, statistical significance was determined by p < 0.05 and Cohen’s *d* of medium and large effect. Data are presented as mean ± SEM.

To determine whether *GBA1* deficiency affects endolysosomal trafficking similarly in neurons and astrocytes, we examined the morphology of early endosomes, recycling endosomes, and lysosomes in *GBA1^+/+^* and *GBA1-*deficient dopaminergic neurons and astrocytes by immunocytochemistry (ICC). *GBA1^+/+^* and *GBA1-*deficient neurons and astrocytes differentiated from 3 different *GBA1^IVS^*^/+^ iPSC clones (#1,2,6), and 3 different control *GBA1^+/+^* clones (#1,2,4) as well as the *GBA1^IVS/IVS^* iPSC clone and a *GBA1^+/+^* clone (#2) treated with 100mM CBE were fixed and co-stained 10-14 days after completing differentiation and maturation with antibodies specific for early endosomes (anti-EEA1), lysosomes (anti-LAMP1), or recycling endosomes (anti-Rab11). Anti-MAP2 and anti-Vimentin were used as markers of neurons and astrocytes, respectively, as both stained the entire cell body reliably, which was necessary for quantitative morphological analysis. To analyze the morphology of early endosomes, lysosomes, or recycling endosomes, we 3D-reconstructed confocal z-stack images of neurons and astrocytes stained for vesicular organelles using Imaris software to determine the volume and size of the stained vesicular structures within the volume of the cell in which those structures were located. The compartment size of early endosomes, lysosomes, and recycling endosomes was then quantified by calculating the sum of the volume of all stained vesicles within a cell divided by the volume of that cell. The diameter of all stained vesicles ascertained in cells from a specific genotype or group was also analyzed, as the size of the vesicles could be altered independent of overall compartment size, indicating impairment in fusion or fission of vesicles. Given the large size of these data sets, we used Cohen’s d to assess the significance between groups in addition to one-way ANOVA to determine significant changes between genotypes.

Representative mages and analyses are shown for one representative *GBA1^+/+^* and *GBA1^IVS/+^* clone however data for all clones is available on Mendeley Data DOI:10.17632/fgswr9ctcr.1.

Morphological analysis of endolysosomal trafficking in *GBA1-*deficient neurons revealed significant alterations compared to control neurons. The diameter of EEA1-stained early endosome vesicles did not differ significantly between *GBA1*-deficient and *GBA1^+/+^* neurons. However, the early endosome compartment of *GBA1^IVS/+^* and *GBA1^IVS/IVS^* neurons was significantly enlarged compared to *GBA1^+/+^*. CBE treatment of *GBA1^+/+^* neurons resulted in the greatest enlargement of the early endosome compartment (Fig 2A-C). EEA1 protein expression was not significantly increased in *GBA1^IVS/+^* neurons compared to *GBA1^+/+^* neurons, but significantly higher in *GBA1^IVS/IVS^* neurons, and the largest increase in EEA1 expression was found in *GBA1^+/+^* neurons treated with CBE (Fig 2D). CBE inhibits not only lysosomal GBA1 but also GBA2, a cytosolic GCase associated with ER and Golgi membranes that can transfer glucose to cholesterol in addition to hydrolyzing glucosylceramide to glucose and ceramide. The increased effect of CBE in *GBA1^+/+^* neurons on early endosomes compared to *GBA1^IVS/IVS^* neurons is likely due to inhibiting both glucocerebrosidase 1 and 2 enzyme function, as *GBA2* has been shown to regulate acidification of the lysosome and endolysosomal trafficking to the Golgi^25^.

Surprisingly, early endosomes were not significantly altered in *GBA1^IVS/+^* or *GBA1^IVS/IVS^* astrocytes compared to *GBA1^+/+^*, either in diameter or compartment size (Fig 2 E-G). CBE treatment of *GBA1^+/+^* astrocytes also did not significantly affect early endosome morphology. Western blot with anti-EEA1 antibody confirmed that EEA1 expression was not significantly altered in *GBA1^IVS/+^, GBA1^IVS/IVS^* and *GBA1^+/+^* astrocytes, as well as *GBA1^+/+^* astrocytes treated with CBE (Fig 2H).

Lysosomes were significantly altered in *GBA1-*deficient neurons and CBE-treated *GBA1^+/+^* astrocytes, but not *GBA1^IVS/+^* or *GBA1^IVS/IVS^* astrocytes. While individual lysosome size was not significantly different between *GBA1^IVS/+^*, *GBA1^IVS/IVS^* and *GBA1^+/+^* neurons treated with CBE compared to *GBA1^+/+^*, total compartment size was enlarged in *GBA1^IVS/+^*, *GBA1^IVS/IVS^* and *GBA1^+/+^* neurons with CBE treatment (Fig 3 A-C). LAMP1 protein expression was significantly increased in *GBA1^+/+^* neurons treated with CBE and *GBA1^IVS/IVS^* neurons but not significantly altered in *GBA1^IVS/+^* compared to *GBA1^+/+^* neurons (Fig 3D). While *GBA1^IVS/+^*, *GBA1^IVS/IVS^* and *GBA1^+/+^* astrocytes treated with CBE did not have significantly enlarged lysosomes, the lysosomal compartment of only *GBA1^+/+^* astrocytes treated with CBE was enlarged compared to *GBA1^+/+^* (Fig 3E-G). However, LAMP1 protein expression was significantly increased in *GBA1^IVS/IVS^* astrocytes and reduced in *GBA1^IVS/+^* astrocytes when compared to *GBA1^+/+^* (Fig 3H).

Recycling endosomes are an important branch point in the endocytic system for trafficking proteins and lipids to the lysosome or the plasma membrane for extracellular secretion within exosomes^15^. Given our prior findings in our *GBA1 mutant Drosophila* model, *dGBA1b,* suggesting that *GBA1* deficiency alters extracellular vesicle biogenesis^12^, we were particularly interested in whether *GBA1* deficiency perturbs recycling endosomes. We investigated whether Rab11, a key regulator and marker of recycling endosomes, was altered in our *GBA1*-deficient and *GBA1^+/+^* neurons and astrocytes. Morphological analysis of RAB11-stained *GBA1^IVS/+^* and *GBA1^IVS/IVS^* neurons did not reveal a significant difference in size of individual recycling endosomes, but the compartment size of recycling endosomes was significantly enlarged in *GBA1^IVS/+^*, *GBA1^IVS/IVS^*, and *GBA1^+/+^* neurons treated with CBE (Fig 4A-C). Western blot of RAB11 protein levels in *GBA1*-deficient neurons revealed a dose-dependent increase in RAB11 protein levels in *GBA1^IVS/+^* and *GBA1^IVS/IVS^* neurons, and even further increase in *GBA1^+/+^* neurons treated with CBE (Fig 4D). *GBA1-*deficient astrocytes again differed from neurons, showing no enlargement of the compartment size nor individual size of recycling endosomes compared to *GBA1^+/+^* (Fig 4E-G). RAB11 protein level was not significantly altered between *GBA1^IVS/+^*, *GBA1^IVS/IVS^* and *GBA1^+/+^* astrocytes treated with CBE (Fig 4H).

To test whether the observed differences in endolysosomal trafficking in *GBA1*-deficient neurons and astrocytes correlate with presence of pathogenic a-synuclein, a hallmark finding in α-synucleinopathies, we examined whether phosphorylated and aggregated a-synuclein was present in our *GBA1-*deficient iPSC-neurons and astrocytes. Neurons stained with an antibody specific to aggregated α-synuclein revealed increased staining in *GBA1^IVS/+^*, *GBA1^IVS/IVS^* and *GBA1^+/+^* neurons treated with CBE (Fig 5A, B). Western blot for phosphorylated α-synuclein at Serine 129 was undetectable in *GBA1^+/+^* neurons, but present in *GBA1^IVS/+^*, *GBA1^IVS/IVS^* and *GBA1^+/+^* neurons treated with CBE (Fig 5C). Western blot using an antibody that recognizes both α-synuclein monomers and oligomers (BD Biosciences #610787) revealed ∼120 kD octamers of α-synuclein in the Triton X-100 insoluble protein fraction of *GBA1^IVS/+^*, *GBA1^IVS/IVS^* and CBE-treated *GBA1^+/+^* neurons, but not in *GBA1^+/+^* neurons (Fig 5D). ICC of astrocytes with an antibody specific to aggregated α-synuclein (Abcam ab209538) revealed an increase in staining only in the *GBA1^IVS/IVS^* astrocytes compared to controls (Fig 5E-F). Pathogenic phosphorylated α-synuclein and α-synuclein oligomers were not significantly increased in *GBA1^IVS/+^*, *GBA1^IVS/IVS^* and *GBA1^+/+^* astrocytes treated with CBE compared to *GBA1^+/+^* astrocytes (Fig 5G, H).

To obtain more global insight into how *GBA1* deficiency alters neurons differentially from astrocytes, we submitted *GBA1^IVS/+^*, *GBA1^IVS/IVS^*and *GBA1^+/+^* neurons and astrocytes for RNA-seq analysis (Fig 6A). *GBA1* expression was not significantly reduced in *GBA1^IVS/+^* neurons or astrocytes compared to *GBA1^+/+^* neurons or astrocytes, respectively. However, *GBA1* expression was downregulated by 80% in *GBA1^IVS/IVS^* neurons compared to *GBA1^+/+^* neurons and 64% in *GBA1^IVS/IVS^* astrocytes compared to *GBA1^+/+^* astrocytes. These reductions likely underestimate the true difference due to transcripts from the *GBA1* pseudogene *GBAP1*, which shares 96% sequence identity with *GBA1*^26^. Interestingly, *GBA2* was upregulated by 103% in *GBA1^IVS/IVS^* astrocytes compared to *GBA1^+/+^* astrocytes but not differentially expressed in *GBA1*-deficient neurons. Of note, *Synuclein-alpha* (*SNCA*) was not differentially expressed in *GBA1^IVS/+^* or *GBA1^IVS/IVS^* neurons or astrocytes compared to control neurons or astrocytes. *EEA1* was downregulated in *GBA1^IVS/+^* neurons by 62% but not astrocytes, and multiple Rab proteins and other proteins involved in macroautophagy and retromer function, including *LRRK2*, were differentially expressed in *GBA1^IVS/+^* neurons compared to *GBA1^+/+^* neurons, but not *GBA1^IVS/+^* astrocytes compared to *GBA1^+/+^* astrocytes. The differentially regulated genes in *GBA1^IVS/+^* neurons and astrocytes compared to *GBA1^+/+^* neurons and astrocytes, respectively, are graphically summarized by volcano plots in Fig 6B-C.

**Figure 6.**
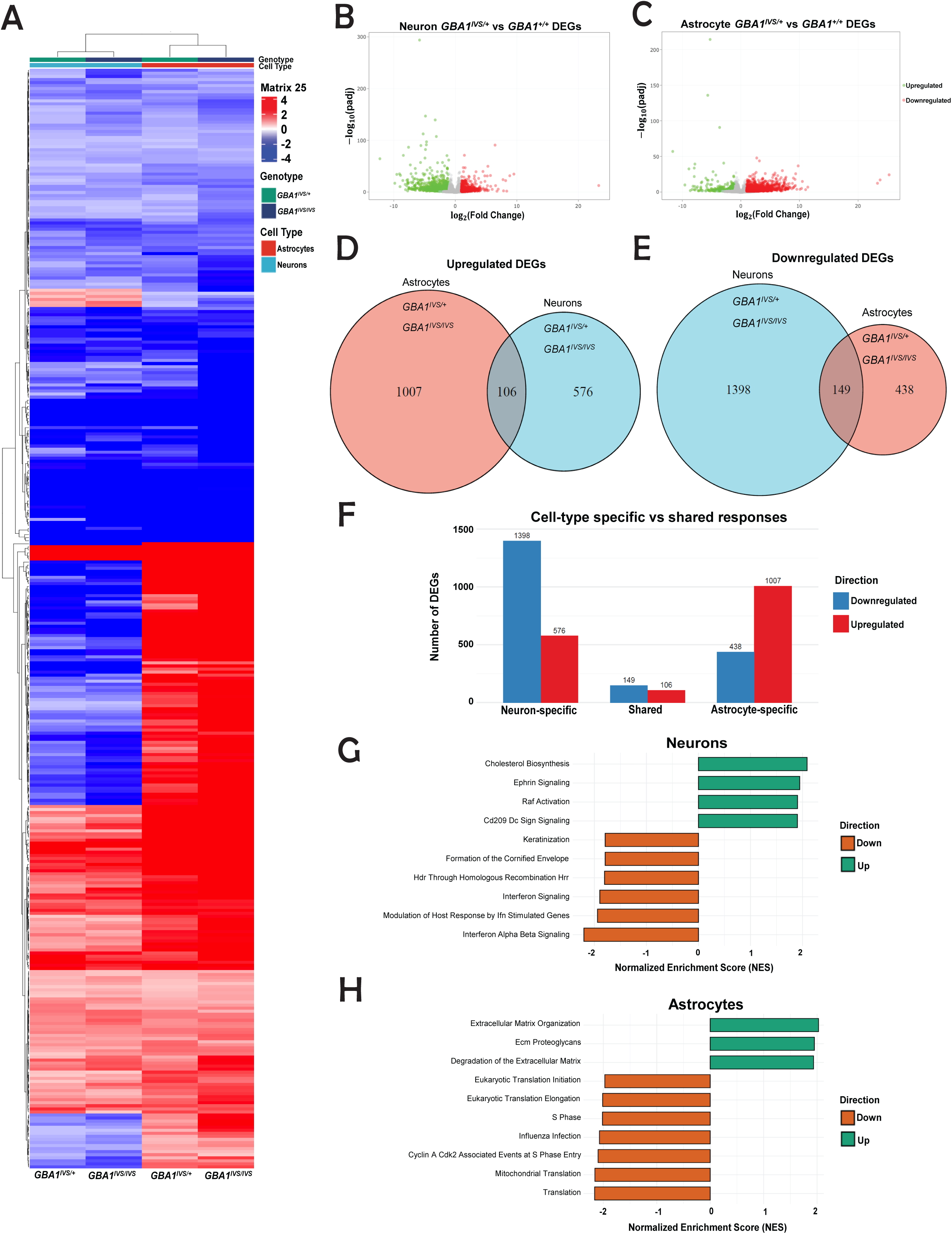
Transcriptomic profiles of *GBA1*-deficient neurons differ significantly from *GBA1*-deficient astrocytes. (**A**) Heat map of all differentially expressed genes with at least log0.5 fold change in *GBA1^IVS/IVS^* and *GBA1^IVS/+^* neurons compared to *GBA1^+/+^* neurons and *GBA1^IVS/IVS^* and *GBA1^IVS/+^* astrocytes compared to *GBA1^+/+^* astrocytes. (**B**) Volcano plot of global transcriptional changes across *GBA1^IVS/+^* versus *GBA1^+/+^* neurons. The log2 fold change of each gene is represented on the x-axis, and the log10 of its adjusted p-value is on the y-axis. Genes with an adjusted p-value < 0.05 and a log2 fold change > 1 are indicated by red dots representing up-regulated genes. Genes with an adjusted p-value less than 0.05 and a log2 fold change less than –1 are indicated by green dots representing down-regulated genes. (**C**) Volcano plot of global transcriptional changes across *GBA1^IVS/+^* versus *GBA1^+/+^* astrocytes. (**D**) Venn diagram of consensus upregulated differentially expressed genes (DEGs) in *GBA1^IVS/+^* and *GBA1^IVS/IVS^* neurons compared to *GBA1^+/+^*, and consensus down-regulated DEGs in *GBA1^IVS/+^* and *GBA1^IVS/IVS^* astrocytes compared to *GBA1^+/+^*. (**E**) Venn diagram of consensus down-regulated differentially expressed genes (DEGs) in *GBA1^IVS/+^* and *GBA1^IVS/IVS^* neurons compared to *GBA1^+/+^*, and consensus down-regulated DEGs in *GBA1^IVS/+^* and *GBA1^IVS/IVS^* astrocytes compared to *GBA1^+/+^*. **(F)** Bar graph of DEGs share between GBA1-deficient neurons and astrocytes and specific to each cell type. **(G)** Gene Set Enrichment Analysis of Reactome pathways using differential gene expression in *GBA1*-deficient neurons relative to control neurons and **(H)** *GBA1*-deficient astrocytes relative to control astrocytes. Expression changes from both *GBA1^IVS/IVS^*and *GBA1^IVS/+^* cells were integrated and ranked by Z-score (Fisher’s method), with the top 10 Reactome pathways by absolute normalized enrichment score (NES) shown. Positive values (green) represent pathways enriched in upregulated genes, negative values (orange) represent pathways enriched in downregulated genes.

To focus on transcriptional changes specific to *GBA1* deficiency, we compared differentially expressed genes (DEGs) in *GBA1^IVS/+^* and *GBA1^IVS/IVS^* neurons compared to *GBA1^+/+^* neurons and found that there was significant overlap in DEGs by log 0.5 fold change or greater, with the majority of DEGs downregulated (1,398 genes) in *GBA1*-deficient neurons and 576 upregulated (Fig 6D-F). In contrast, the majority of DEGs in *GBA1*-deficient astrocytes were upregulated (1,007) compared to downregulated (438) (Fig 6D-F). DEGs common to both *GBA1^IVS/+^* and *GBA1^IVS/IVS^* neurons or astrocytes at a threshold of log0.5 fold or greater were termed “consensus DEGs.” We then compared neuron and astrocyte consensus DEGs and found that the majority of DEGs were non-overlapping, supporting our *in vitro* findings that *GBA1* deficiency has a cell type-specific effect (Figure 6D-F).

We performed Gene Set Enrichment Analysis (GSEA)^27^ using the Reactome pathway database to compare DEGs specific to *GBA1-*deficient neurons versus those specific to astrocytes, and found strikingly different cellular pathways were affected (Figure 6G-H). *GBA1*-deficient neurons upregulated genes involved in cholesterol biosynthesis, axon guidance, Raf activation, and immune cell signaling, whereas downregulated genes in *GBA1*-deficient neurons were primarily implicated in immune signaling (Fig 6G). GSEA for Reactome pathways in *GBA1*-deficient astrocytes revealed upregulation of genes involved in extracellular matrix biogenesis, while downregulated genes in *GBA1*-deficient astrocytes included genes involved in translation (Fig 6H), suggesting a more reactive astrocyte population promoting extracellular remodeling leading to gliosis^28^.

We examined in greater depth the 255 DEGs shared by both *GBA1*-deficient neurons and astrocytes to gain insight into common pathways regulated by *GBA1* in both cell types. Overall, *GBA1*-deficient astrocytes tended to have greater change in differential expression of concordant genes compared to *GBA1*-deficient neurons, as indicated in a heatmap of the average expression level between *GBA1^IVS/+^* and *GBA1^IVS/IVS^* neurons, and *GBA1^IVS/+^* and *GBA1^IVS/IVS^* astrocytes (Fig 7A). The concordance is also demonstrated in a scatterplot log2 fold change DEGs in *GBA1*-deficient astrocytes versus log2 fold change DEGs in *GBA1*-deficient neurons, where Pearson r = 0.889 (Fig 7B). DEGs shared by *GBA1*-deficient neurons and astrocytes included genes involved in lipid metabolism, lysosome and autophagy function, mitochondrial function, ER stress response, ER-membrane contacts and epigenetic pathways (Fig 7C-E). Perturbations of these pathways, particularly in sphingolipid metabolism, ER membrane interactions, lysosomes and autophagy were anticipated given glucocerebrosidase is a lysosomal enzyme that catabolizes the glycolipid glucocerebroside into ceramide. In addition to altering sphingolipid abundance, which would likely impact membrane phospholipid composition, ceramides can also modulate lipid rafts, membrane fluidity, and exocytosis^29–31^.

**Figure 7.**
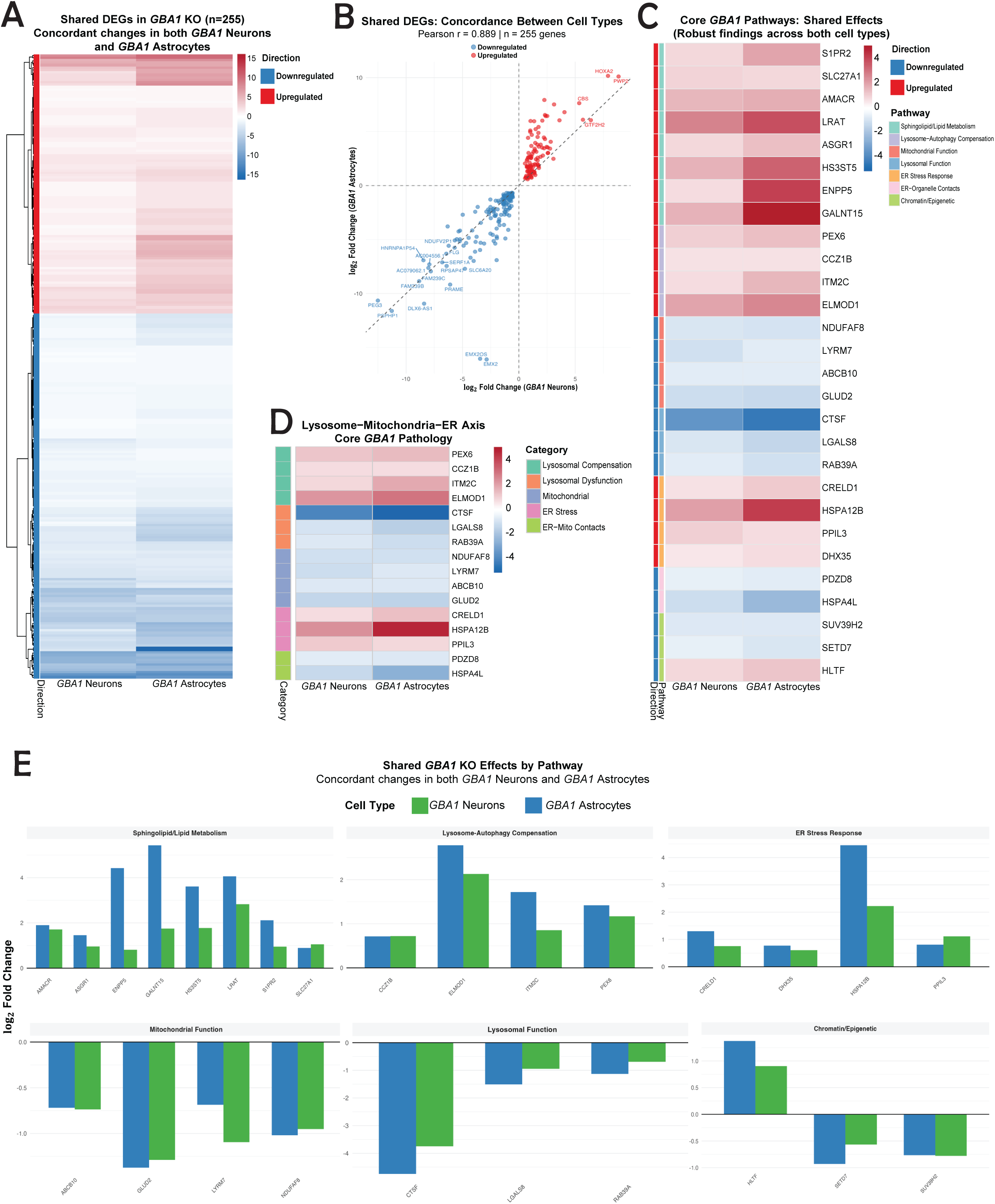
Transcriptomic profile of differentially expressed genes common to *GBA1*-deficient neurons and astrocytes. (**A**) Heat map of the 255 differentially expressed genes common to both GBA1-deficient neurons and astrocytes. The z-score for each differentially expressed gene with at least log 0.5 fold change in *GBA1^IVS/+^* and *GBA1^IVS/IVS^* compared to *GBA1^+/+^* was averaged for neurons and astrocytes. **(B)** Scatter plot of differentially expressed genes between *GBA1*-deficient neurons and astrocytes reveals a linear relationship with Pearson r = 0.889. **(C)** Heatmap of concordant differentially expressed genes in specific cellular pathways between *GBA1*-deficient neurons and astrocytes. **(D)** Heatmap of concordant differentially expressed genes within the lysosome-mitochondria-ER axis between *GBA1*-deficient neurons and astrocytes. **(E)** Bar graph of concordant differentially expressed genes in *GBA1*-deficient neurons and astrocytes within various cellular pathways.

Most of the DEGs in *GBA1*-deficient astrocytes and neurons are not overlapping, suggesting cell-type specific perturbation of pathways. We examined the neuron-specific and astrocyte-specific consensus DEGs to determine which cell type-specific pathways might be altered by *GBA1* deficiency. We found that lipid metabolism genes, including APOE and CH25H, were differentially upregulated in *GBA1*-deficient astrocytes (Fig 8A, C). *GBA1*-deficient astrocytes differentially upregulated genes associated with reactive astrocytes and gliosis, including *CXCL1* and *CHI3L1* suggestive of reactive neurotoxic A1 astrocytes^32^(Fig 8A-B), while interferon response genes were differentially downregulated in *GBA1*-deficient neurons compared to astrocytes (Fig 8F). *GBA1*-deficient astrocytes also differentially upregulated cholesterol efflux and sphingolipid genes, while *GBA1*-deficient neurons differentially upregulated genes involved in cholesterol synthesis and fatty acid desaturation (Fig 8C).

**Figure 8.**
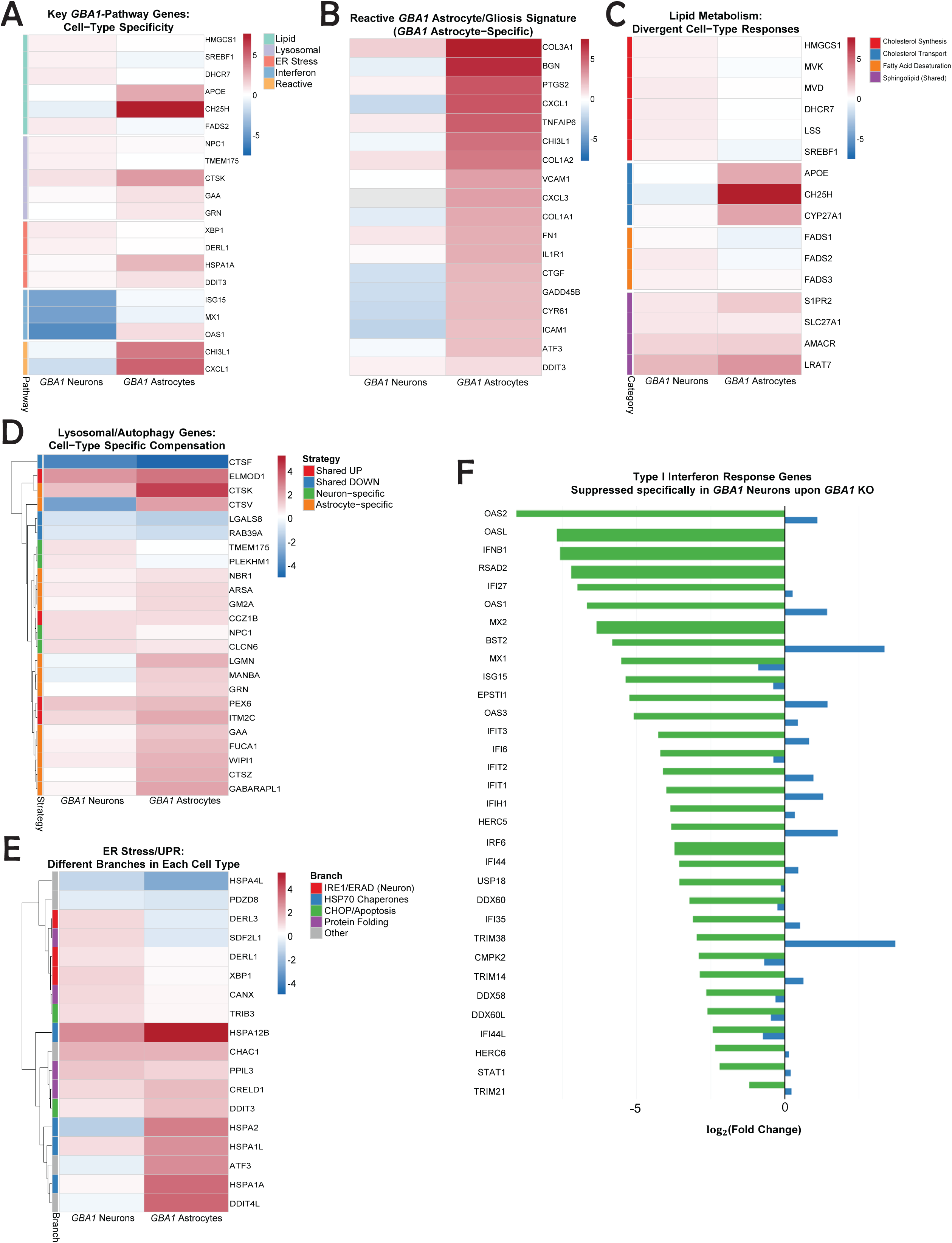
Transcriptomic profile of differentially expressed genes specific to *GBA1*-deficient neurons and astrocytes. (**A**) Cell type specific heat map of mean z-scores for differentially expressed genes in *GBA1*-deficient neurons and astrocytes involved in lipid, lysosomal, ER stress, interferon, and reactive pathways. (**B**) Cell type-specific heat map of differentially expressed genes due to *GBA1*-deficiency involved in reactivity and gliosis. (**C**) Cell type-specific heat map of differentially expressed genes in *GBA1*-deficient neurons and astrocytes involved in lipid metabolism. (**D**) Cell type-specific heat map of differentially expressed genes in *GBA1*-deficient neurons and astrocytes involved in autophagy and lysosome pathways. (**E**) Cell type-specific heat map of differentially expressed genes in *GBA1*-deficient neurons and astrocytes involved in ER stress and unfolded protein response pathways. **(F)** Type I interferon response genes differentially regulated in *GBA1*-deficient neurons.

Surprisingly, *EEA1* mRNA expression was reduced in *GBA1^IVS/+^* neurons by 63% compared to *GBA1^+/+^* neurons. *EEA1* was not differentially expressed in *GBA1^IVS/+^* astrocytes compared to *GBA1^+/+^* astrocytes, however several other genes involved in intracellular membrane trafficking and autophagy (Fig 8D) and the ER stress/UPR response (Fig 8E) were upregulated in *GBA1*-deficient astrocytes. *GBA1*-deficient neurons displayed significant downregulation of genes responsive to type 1 interferon (Fig 8F). Overall, the transcriptomic profiles of *GBA1*-deficient neurons were divergent from *GBA1*-deficient astrocytes, with downregulation of inflammatory pathways, and upregulation in cholesterol synthesis, whereas *GBA1*-deficiency in astrocytes specifically upregulated APOE and lipid efflux as well as extracellular matrix biogenesis pathways and downregulated genes involved in translation.

## Discussion

The underlying cellular mechanisms responsible for the association between *GBA1* deficiency and both PD/DLB risk and faster disease progression have been difficult to elucidate. Our study using human iPSC-derived neurons and astrocytes reveals a cell type specific differential effect of *GBA1* deficiency, where neurons are more susceptible to perturbation in endolysosomal trafficking and α-synucleinopathy compared to astrocytes. We found that early endosomes, lysosomes, and recycling endosomes were all enlarged in cellular compartment size in *GBA1*-deficient neurons. However, only treatment with CBE of *GBA1^+/+^* astrocytes resulted in enlarged lysosome cellular compartment size, while *GBA1*-deficient astrocytes did not significantly differ in early or recycling endosome or lysosome compartment size compared to *GBA1^+/+^* astrocytes.

These differences correlate with the presence of pathogenic α-synuclein aggregation and phosphorylation in neurons but not astrocytes. Transcriptomic profiling of *GBA1-*deficient neurons and astrocytes confirmed cell-type specific effects, where the majority of DEGs in *GBA1*-deficient neurons were downregulated, while the majority of DEGs in *GBA1*-deficient astrocytes were upregulated. Upregulated DEGs in *GBA1*-deficient neurons included genes in cholesterol synthesis and lipid metabolism pathways, and downregulation of genes involved in inflammatory pathways. In contrast, *GBA1*-deficient astrocytes had upregulation of genes involved in extracellular matrix biogenesis, neurotoxic astrocyte reactivity, and downregulation of genes involved in translational machinery. These findings suggest that dopaminergic neurons are more susceptible to endolysosomal trafficking defects due to *GBA1*-deficieny, leading to α-synucleinopathy. However, *GBA1*-deficiency may have a deleterious impact on astrocytes that is independent of α-synucleinopathy.

Prior work has found enlarged lysosomes and autophagolysosomes in multiple models of *GBA1-*deficiency^33,34^ and human postmortem tissue^35^. However, early endosome defects have not been characterized in *GBA1-*deficient neurons. Both *GBA1^IVS/+^* and *GBA1^IVS/IVS^* neurons were found to have an enlarged early endosome compartment size and increased protein expression of EEA1 by Western blot, even though EEA1 mRNA expression was reduced in *GBA1^IVS/+^* neurons compared to *GBA1^+/+^* neurons. These findings suggest that *GBA1* deficiency reduces the fission and/or fusion of early endosomes, resulting in accumulation of enlarged early endosomes and EEA1 protein. Similar alterations in lysosomes and recycling endosomes were found in *GBA1^IVS/+^* and *GBA1^IVS/IVS^* neurons. Surprisingly, we did not observe enlarged compartment size of early endosomes, lysosomes or recycling endosomes in *GBA1^IVS/+^*, *GBA1^IVS/IVS^* or *GBA1^+/+^* astrocytes treated with the GCase inhibitor CBE. Together, our findings indicate that endolysosomal trafficking in neurons is more susceptible to *GBA1* deficiency than astrocytes, although when GCase activity is severely reduced or inhibited, endolysosomal trafficking is also impaired in astrocytes.

The increased susceptibility of neurons to effects of *GBA1* deficiency on endolysosomal trafficking is correlated with the presence of pathogenic α-synuclein phosphorylation and aggregation. Human postmortem tissue and *in vitro* models suggest that phosphorylation of Ser129 may promote pathogenic α-synuclein aggregation^36,37^. Recent work revealed phosphorylation of Ser129 impairs interactions between the ER and mitochondrial membranes, leading to mitochondrial dysfunction and calcium dysregulation^38^. Interestingly, pathogenic forms of α-synuclein were detected in our *GBA1*-deficient neurons but not astrocytes and transcriptomic analysis of our *GBA1*-deficient neurons revealed upregulation in genes involved in ER-mitochondrial membrane interactions. Further elucidation of how *GBA1* deficiency regulates phosphorylation of Ser129 of α-synuclein and perturbs mitochondrial-associated membrane interactions could provide insight into the pathogenesis of α-synucleinopathies.

Several studies have suggested that astrocytes may have more efficient endolysosomal trafficking compared to neurons, which could be protective due to increased astrocytic endocytosis of extracellular pathologic aggregating proteins that may otherwise seed pathology in neurons^21,22,39^. Astrocytes differentiated from human iPSCs were found to more rapidly internalize extracellular α-synuclein compared to iPSC-dopaminergic neurons. When astrocytes were co-cultured with neurons with biallelic mutations in *ATP13A2*, which cause a recessive form of Parkinsonism, there was reduced neuronal α-synuclein, although phosphorylated and aggregated forms of α-synuclein were not examined in this study^22^. In *Drosophila*, *GBA1* was shown to be necessary for astrocyte-mediated neuronal pruning during sleep^40^. These differential cell type-specific effects in *GBA1* function may have a protective role as well, as expression of wildtype *dGBA1b* in *Drosophila* astrocytes was sufficient to prevent protein aggregation in neurons in *GBA1-*deficient flies^40^. Our findings further support a cell type-specific role for *GBA1*, in which neurons have increased susceptibility to endolysosomal trafficking dysfunction and development of α-synucleinopathy.

While most investigations have used models heterozygous for the more common pathogenic *GBA1* variants N370S or L444P, this study examines iPSC-derived neurons and astrocytes carrying the *GBA1* IVS2+1 variant as it is considered to be a variant leading to complete absence of GCase protein, rather than a missense variant that may cause additional cellular effects through protein misfolding and ER stress. The effects that we observed in our *GBA1^IVS^*^/+^ neurons trended towards increased severity in homozygous *GBA1^IVS/IVS^* neurons and *GBA1^+^*^/+^ neurons treated with CBE, supporting reduced GCase function rather than a gain of function etiology for the observed alterations in endolysosomal trafficking and α-synuclein pathology. However, it will be interesting to see if iPSC-derived cells carrying other *GBA1* variants such as N370S and L444P recapitulate our cell-type specific findings.

RNA-seq analysis of our *GBA1*-deficient neurons and astrocytes revealed dramatically different gene expression signatures, supporting our findings that *GBA1* deficiency has cell type-specific effects on divergent cellular pathways. *GBA1*-deficient neurons were found to have upregulation of genes involved in exocytic, synaptic, and mitochondrial membranes and down-regulation of inflammatory genes. In contrast, *GBA1*-deficient astrocytes resulted in downregulation of genes involved in transcription and upregulation of extracellular matrix genes, suggestive of astrocyte reactivity leading to gliosis. Profiling of *GBA1*-deficient astrocytes based on markers for astrocyte subpopulations revealed the upregulation of genes consistent with A1 reactive astrocytes^32^. Neurotoxic A1 astrocytes are implicated in PD pathogenesis through microglial activation, leading to dopaminergic cell death^41^. *GBA1* deficiency has been found to reduce inflammatory response in primary astrocytes from *GBA1* D409V knock in mice^42^, and interestingly, these astrocytes did not have impaired degradation of α-synuclein monomers or fibrils. *In vitro* studies examining the inflammatory response of our *GBA1* IVS2+1 mutant astrocytes and co-culturing with wildtype versus *GBA1*-deficient neurons will elucidate the role of *GBA1* deficient astrocytes in PD pathogenesis. However, our transcriptomic analysis suggests that *GBA1* deficiency may independently promote the transition of astrocytes into the pathogenic A1 state, contributing to pathogenesis independently of endolysosomal trafficking defects and α-synuclein aggregation.

Glucocerebrosidase is an important enzyme for ceramide and glucosphingosine metabolism, and several studies have underscored the influence of lipid metabolism on neurodegeneration. Ceramidase inhibition with carmofur, resulting in increased ceramide level, was found to reverse α-synuclein aggregation in *GBA1-*deficient HEK293T cells^43^. Given that astrocytes are the main cell type in the central nervous system responsible for lipid metabolism and efflux of lipids to neurons^18^, we anticipated that *GBA1* deficiency would have a significant impact on astrocyte lipid metabolism.

Interestingly, there was significant upregulation of genes involved in lipid metabolism pathways, including cholesterol efflux, fatty acid metabolism, inflammatory lipid signaling, and membrane remodeling (Fig 6,8). Alterations in lipid metabolism in both neurons and glia have been implicated in PD, neurodegeneration, and aging^44–46^. Our findings support differential lipid metabolism alterations due to *GBA1* deficiency in neurons and astrocytes, and further characterization of lipid profiles will elucidate how alterations in each cell type influences neurodegeneration.

Our model of PD-*GBA1* deficiency using iPSC differentiated neurons and astrocytes reveals the cell-specific effects and potential neuroprotective roles for astrocyte specific *GBA1*. Importantly, this study underscores the role that glial cells have in PD and how known risk factors like *GBA1* may impair glial function independently of α-synuclein aggregation, greatly contributing to the progression of neurodegeneration. Although *GBA1* deficiency in astrocytes did not result in endolysosomal trafficking defects and α-synucleinopathy as in *GBA1*-deficient neurons, transcriptomic profiling suggests *GBA1*-deficiency causes pathogenic reactive astrocyte changes and extracellular matrix changes. These extracellular matrix changes may represent gliosis^28^. Our findings lay a strong foundation to investigate further mechanisms that underlie *GBA1-*deficient pathology and could reveal novel therapeutic targets for slowing progression of neurodegeneration in a broader population of α-synucleinopathy patients.

## Conclusion

In conclusion, our study examined endolysosomal trafficking in human iPSCs carrying the *GBA1 IVS2+1* null variant differentiated into dopaminergic neurons and astrocytes. In this model, we demonstrated cell-type specific enlargement of the early endosome, lysosome, and recycling endosome compartment in *GBA1*-deficient neurons but not astrocytes. *GBA1*-deficient neurons but not astrocytes were found to have α-synucleinopathy. RNA sequencing reveals cell type-specific effects of *GBA1* deficiency, with significantly different cellular pathways affected in *GBA1*-deficient neurons and *GBA1*-deficient astrocytes. Our model provides insight into the cell-type specific role of *GBA1-*deficiency and will be valuable for further defining the individual contributions of cell types to α-synucleinopathy and neurodegeneration.

## Materials & Methods

### IPSC Generation

Fibroblasts were obtained from a deidentified individual diagnosed with PD and heterozygous for *GBA1^IVS2+1^* ascertained by the Parkinson’s Genetic Research Study (PI Zabetian, VA Puget Sound IRB Protocol #1587534) and were reprogrammed to generate iPSCs by the University of Washington Institute for Stem Cell and Regenerative Medicine Ellison Stem Cell Core. All clones were verified by sequence analysis using the following primers:

### PCR Primers

**F:** GTTGTCACCCATACATGCCC

**R**: CTCTCATGCATTCCAGAGGC

**Product length**: 7,050

### Sequencing Primers

**F:** GTGGGCCTTGTCCTAATGAA

**R:** CAAAGGACTATGAGGCAGAA

All iPSC clones were screened for mycoplasma (Lonza #LT07-118) and chromosomal abnormalities by karyotyping by Diagnostic Cytogenetics Incorporated, Seattle, WA, or whole genome sequencing by Karyostat+ ThermoFisher (S Fig 2).

Generation of isogenic control *GBA1^IVS/IVS^* Rev, *GBA1^IVS/IVS^* and *GBA1^IVS/+^* A4 clones One million iPSCs derived from patients harboring the *IVS2+1 G>A* mutation at the *GBA1* locus were electroporated with *Cas9* (0.3-0.6 µM, Sigma) and gRNA (GGCATCAGATGAGTGAGTCA)(1.5-3 µM, Synthego) as RNP complex along with ssDNA donor:

AGTAGGGTAAGCATCATGGCTGGCAGCCTCACAGGATTGCTTCTACTTCAGGCAGTGTCGTGGGCATCAGgtgagtgagtcaaggcagtggggaggtagcacagagcctcccttctgcctcatagtcctttggtagccttc (2uM, IDT) using Amaxa nucleofector (Human Stem Cell kit 2) in presence of Rho-associated, coiled-coil containing protein kinase (ROCK) inhibitor. Individual colonies were hand-picked and plated into 96 well plates. DNA was extracted using Quick Extract DNA extraction solution (Epicentre#QE09050) and nested PCR was performed with the following primers:

**F1:** GGCTTGCTTTTCAGTCATTCCTC

**R1**: ACATGGAGAATGGACACATCTGCTA

**Product length**: 635

The PCR product was purified using EXO-SAP enzyme (ThermoFisher) and sent for Sanger sequencing analysis (through Genewiz). The integrity of the *GBA1* pseudogene, *GBAP1* was also assessed by nested PCR and Sanger sequencing using the following primers:

**F1:** CTTCGGGTAGGGTAAGCATCAT

**R1**: CTGACAAGGATGTGGAGAACGGA

**Product length**: 283

See Supplemental Figure 1 for further details.

### Differentiation of iPSCs into NPCs

Differentiation of iPSC into neural progenitor cells (NPCs) was performed using STEMCELL Technologies reagents and procedures specified in the Technical Manual of the STEMdiff^TM^ SMADi Neural Induction Kit (Cat No. 08581). Briefly, a single cell suspension of iPSCs was seeded onto Matrigel-coated plates at a density of 2.5-3×10^5^ cells/cm^2^ in STEMdiff^TM^ Neural Induction Medium +SMADi. Cells were passaged every 6-9 days and replated at a density of 1.5-2×10^5^ cells/cm^2^. After three weeks in STEMdiff^TM^ Neural Induction Medium + SMADi, a small portion of the cells was fixed and stained with the following antibodies to evaluate presence of NPC markers: mouse anti-Nestin (Santa Cruz # sc-23927, 1:50), rabbit anti-Sox1 (Cell Signaling #4194S, 1:100), and anti-Pax6 (Santa Cruz # sc-81649, 1:50). Verified NPCs were frozen down or expanded for neuronal or astrocyte differentiation.

### Midbrain Dopaminergic Neuron Differentiation

NPCs passaged less than 5 times were differentiated into dopaminergic neurons using the reagents and procedures specified in the Technical Manual of the STEMdiff^TM^ Midbrain Neuron Differentiation Kit (STEMCELL Technologies Cat No.100-0038). NPCs were seeded onto Matrigel-coated 100mm dishes at a density of 2-2.5×10^5^ cells/cm^2^ and grown in Midbrain Neuron Differentiation Medium + Supplement for 1 week or until neural rosettes were formed, then passaged and replated to a density of 6-8×10^4^ cells/cm^2^ and grown in STEMdiff^TM^ Midbrain Neuron Maturation Medium + Supplement for 2 weeks.

### Astrocyte Differentiation

NPCs were differentiated into astrocytes using the reagents and procedures specified in the Technical Manual of the STEMdiff^TM^ Astrocyte Differentiation Kit (STEMCELL Technologies Cat No. 100-0013). NPCs were seeded onto Matrigel-coated 100mm dishes at a density of 1.5-2×10^5^ cells/cm^2^ and grown in STEMdiff^TM^ Astrocyte Differentiation Medium + Supplement for 3 weeks with passaging every 7-10 days. Astrocyte progenitor cells (APCs) were then seeded onto Matrigel-coated 100mm dishes at a density of 1.5-2×10^5^ cells/cm^2^ and grown in STEMdiff^TM^ Astrocyte Maturation Medium + Supplements A&B for 2 weeks with passaging every 7-10 days. After 2 weeks in Astrocyte Maturation Medium, a small portion of cells was fixed and stained to evaluate the presence of mature astrocyte markers: mouse anti-glial fibrillary acidic protein (GFAP) (Sigma-Aldrich G3893, 1:400), mouse anti-Vimentin (Invitrogen #MA5-11883, 1:500), rabbit anti-Vimentin (Cell Signaling Technology 5741S, 1:100), rabbit anti-S100β (Proteintech 15146-1-AP, 1:100), and rabbit anti-Aquaporin 4 (AQP4Invitrogen #PA5-85767, 1:100).

### GCase Enzyme Activity Assay

iPSC-derived neural progenitor cell (NPC) pellets were lysed with lysis buffer consisting of 150 mM citrate phosphate buffer, 0.25% taurocholic acid sodium salt hydrate (Thermo Scientific A18346.03), and 0.25% Triton X-100 (Thermo Scientific A16046-AE) for 30 minutes at 4°C. Lysates were homogenized via sonication and spun down at 14000 rpm at 4°C for 15 minutes; supernatant was then collected. The lysates were added in triplicate to a 96-well plate at 2.5, 5, and, 10 µL with 40 µL assay buffer, consisting of lysis buffer plus 3% BSA (Proliant Biologics 9048-46-8), and 4 mM 4-methylumbelliferyl-β-D-glucopyranoside (Cayman 20948). The plate was then incubated at 37°C, protected from light. After 1 hour, the assay was halted with 100 µL of 0.2 M glycine (Fisher Scientific G46-500). Absorbance at 450 nm was measured with a SpectraMax iD3 plate reader (Molecular Devices, San Jose, CA).

### CBE inhibition of neurons and astrocytes

*GBA1^+/+^* clone #2 cells were treated with conduritol B epoxide (CBE), an inhibitor of GCase activity. After reaching day 0 of maturation, a portion of healthy control neurons and astrocytes was treated with 100mM CBE (Enzo Life Sciences BML-S104-0025) that was replenished every 2-3 days with media changes at 37°C. CBE inhibition was maintained until D10, whereupon the cells were fixed and/or pelleted.

### Immunocytochemistry

iPSC-derived neurons and astrocytes grown on coverslips were fixed in 4% paraformaldehyde and were blocked and permeabilized by incubating in blocking buffer (2.5% BSA, 0.1% Triton X-100 in PBS) for 30 minutes. The fixed cells were incubated overnight at 4°C with primary antibodies dissolved in blocking buffer and were incubated at room temperature for 2 hours with secondary antibodies dissolved in blocking buffer. The following primary antibodies were used: mouse anti-α-synuclein aggregate (Abcam ab209538, 1:250), mouse anti-EEA1 (BD Biosciences #610456, 1:500), rabbit anti-LAMP1 (Cell Signaling #9091S, 1:100), rabbit anti-Rab11 (Abcam #ab3612, 1:1000), mouse anti-MAP2 (ThermoFisher #131500, 1:250), rabbit anti-MAP2 (Cell Signaling Technology #4542S, 1:500), mouse anti-tyrosine hydroxylase (TH) (Santa Cruz #sc-25269), mouse anti-Vimentin (Invitrogen #MA5-11883, 1:500), and rabbit anti-Vimentin (Cell Signaling Technology 5741S, 1:100).

The following secondary antibodies were used: AlexaFluor 488 goat anti-mouse (ThermoFisher A1101, 1:500), AlexaFluor 488 goat anti-rabbit (ThermoFisher A11008, 1:500), AlexaFluor 594 goat anti-mouse (ThermoFisher A32742, 1:500), AlexaFluor 594 goat anti-rabbit (ThermoFisher A32740, 1:500), and DAPI (Santa Cruz Biotechnology sc-3598, 1:1000).

### Confocal Microscopy and Imaris Volume Analysis

Coverslips of fixed and stained neurons or astrocytes were mounted on glass slides with ProLong Gold^TM^ Antifade Mountant (ThermoFisher P10144). Confocal images were obtained using a Nikon A1R Confocal imaging system with Nikon Elements software.

For each marker and genotype pair, at least 20 images were obtained at 60x magnification with an oil-immersion lens using Nikon immersion oil type F. The Nyquist window was 1 µm and the Z-stack was 2 µm in total centered on the nucleus with 0.1 µm steps. Confocal microscopy images were converted to.ims files and subsequently deconvolved using uniform parameters across all images. Using IMARIS 9.9.0, a 3D imaging and analysis software, three fluorescence signals, TRITC, FITC, and DAPI, representing puncta, cell bodies, and nuclei, respectively, were reconstructed into 3D surfaces, using similar methods from prior published studies^47,48^. The DAPI and FITC surfaces were merged to generate a combined representation of the cell. This merged surface, representing the cell of interest, was then segmented to isolate a single, intact cell body. TRITC surfaces were filtered based on their overlap with the cell of interest. Volumetric ratio data, puncta size, and mean intensity values of TRITC surfaces were collected and analyzed using GraphPad Prism 10. At least 15 cells were analyzed per replicate, three different clones per genotype. Resulting data was analyzed by one way ANOVA for normally distributed data and Wilcoxon rank sum test with Bonferroni correction for multiple comparisons or Student’s t-test if comparing only two groups.

Effect size based on Cohen’s d as well as p value were determined using GraphPad Prism 10.

### Western Blotting

For soluble proteins, lysates of iPSC-neurons or astrocytes were obtained using RIPA buffer with Halt protease and phosphatase inhibitor cocktail (ThermoScientific #78441). For insoluble proteins such as high molecular weight α-synuclein aggregates and phospho-Serine129 α-synuclein, cells were extracted in 1% Triton X-100 lysis buffer (150 mM NaCl, 50 mM Tris-HCl, 1 mM EDTA). Cells were sonicated 3 times at 60% amplitude for 10 seconds on and 5 seconds off, then centrifuged at 4°C 15,000xg for 20 minutes. The supernatant was collected for Triton X-100 lysis soluble proteins and the remaining pellet was resuspended in 2% SDS in DI H2O with protease inhibitor cocktail (Sigma-Aldrich p8340-5ML) for Triton X-100 lysis insoluble proteins. Cells were sonicated 3 times at 60% amplitude for 10 seconds on and 5 seconds off, then centrifuged at 4°C 15,600xg for 10 minutes, then boiled for 10 minutes. Approximately 20 ug of protein was loaded from each sample and run on 4-20% Tris-glycine gels, then transferred to PVDF membrane using an iBlot system. Membranes were probed for the following primary antibodies: rabbit anti-glucocerebrosidase (Millipore Sigma #G4171, 1:250), mouse anti-EEA1 (BD Biosciences #610456, 1:2500), rabbit anti-LAMP1 (Cell Signaling #9091S, 1:1000), rabbit anti-Rab11 (Abcam #ab3612, 1:2000), mouse anti-actin (EMD Millipore MAB1501, 1:8000), and secondary goat anti-mouse HRP (Bio-Rad 1706516, 1:1000-7500) or goat anti-rabbit HRP (Bio-Rad #1721019, 1:2000-10000) and visualized with Pico ECL (ThermoFisher #34580) or Femto ECL (ThermoFisher #34096).

For detecting α-synuclein high molecular weight oligomers, Triton X-100 lysis insoluble fractions were run on a 4-20% Tris-Glycine gel and transferred onto PVDF membrane using the iBlot system. The membrane was blocked in 5% non-fat milk in PBST for 30 minutes, then fixed with 0.4% paraformaldehyde for 30 minutes.

Membranes were incubated with primary mouse anti-α-synuclein antibody (BD Biosciences #610787, 1:250) overnight at 4°C in blocking buffer, respectively, then incubated with goat anti-mouse HRP (BioRad #170-6516, 1:1000) and visualized with Pico ECL (ThermoFisher #34580).

For detecting phospho-Serine129 α-synuclein, Triton X-100 lysis insoluble fractions were run on a 4-20% Tris-Glycine gel and transferred onto PVDF membrane using the iBlot system. The membrane was blocked with 0.4% paraformaldehyde in TBS for 30 minutes, then blocked in blocking buffer 5% BSA (Sigma-Aldrich A9418) in TBST for 30 minutes. Membranes were incubated with primary rabbit anti phospho-Serine129 α-synuclein antibody (abcam ab51253, 1:1000) overnight at 4°C in blocking buffer, then incubated with goat anti-rabbit HRP (BioRad #1721019, 1:2000) and visualized with Pico ECL (ThermoFisher #34580).

At least three independent experiments were performed with three biological replicates of each genotype and antibody. Signal was normalized to actin within each Western blot for loading control. Normalized western blot data were log-transformed when necessary to stabilize variance before means were compared using one-way ANOVA.

### RNA Sequencing

Three replicates of the following iPSC clones differentiated into astrocytes and midbrain dopaminergic neurons were submitted to Azenta Life Sciences for RNA sequencing: *GBA1^IVS^*^/+^ clones #1,2,6, control *GBA1^+/+^* clones #1,2,3, *GBA1^+/+^ Rev* isogenic control clone, and a matched control *GBA1^IVS^*^/+^ A4 and homozygous null *GBA1^IVS/IVS^.* Cell pellets of 1×10^6^ were submitted to Azenta for RNA extraction, cDNA libraries were preparation using polyA capture, and sequencing was performed on an Illumina NovaSeq X Plus using 2×150bp paired-end reads at a final read depth of ∼20 million reads per sample. Raw data was processed by Azenta’s standard bioinformatics analysis workflow. Briefly, reads were checked for quality control and adapter sequences were trimmed using Trimmomatic v.0.36. Sequence data was then aligned to the GRCh38 Homo Sapiens reference genome using STAR aligner v.2.5.2b. The resulting BAMs were used to generate hit counts using Subread v.1.5.2, with only exonic regions included for downstream RNA-seq analyses. The RNAseq data analyzed in this publication have been deposited in NCBI’s Gene Expression Omnibus^49^ and are accessible through GEO Series accession number GSE315738 (https://www.ncbi.nlm.nih.gov/geo/query/acc.cgi?acc=GSE315738).

Differential gene expression using DESeq2 was used to compare *GBA1^+^*^/+^ neurons with *GBA1^+^*^/+^ astrocytes, *GBA1^+^*^/+^ neurons with *GBA1^IVS^*^/+^ neurons, *GBA1^+^*^/+^ neurons with *GBA1^IVS/IVS^* neurons, *GBA1^+^*^/+^ astrocytes with *GBA1^IVS^*^/+^ astrocytes, *GBA1^+^*^/+^ astrocytes with *GBA1^IVS/IVS^* astrocytes, and *GBA1^IVS^*^/+^ neurons with *GBA1^IVS^*^/+^ astrocytes. *GBA1^IVS^*^/+^ neurons were also compared to *GBA1^IVS^*^/+^ A4 neurons, and *GBA1^IVS^*^/+^ astrocytes were compared to *GBA1^IVS^*^/+^ A4 astrocytes. *GBA1^+^*^/+^ neurons or astrocytes were compared to isogenic *GBA1^+^*^/+^ Rev neurons or astrocytes to determine whether the isogenic controls were significantly different from the unrelated age-and sex-matched healthy control cell line. p-values and log2 fold changes were calculated using the Wald test, with cutoffs of adjusted p-value < 0.05 and absolute log_2_ fold change >0.5. Significant genes were identified as differentially expressed relative to their respective controls. Genes identified as DEGs using these cutoffs were compared for *GBA1^IVS^*^/+^ and *GBA1^IVS/IVS^* for each cell type (astrocytes and neurons); those meeting the threshold for DEGs in both conditions and expressed in the same direction (upregulated or downregulated) were considered as consensus DEGs. Venn diagrams were created using the R package VennDiagram 1.7.3. Heatmaps of gene expression data were generated using the package pheatmap 1.0.13 using log_2_ fold changes calculated by DESeq2.

For gene set enrichment analysis (GSEA), we identified genes and pathways consistently dysregulated across *GBA1* mutant conditions by combining differential expression results from *GBA1^IVS^*^/+^ and *GBA1^IVS/IVS^* cells using Fisher’s combined statistic. Differential expression tables from DESeq2 for *GBA1^IVS^*^/+^ and *GBA1^IVS/IVS^* cells were compared, and any gene absent from either dataset was dropped for these analyses. For genes present in both comparisons, we used the Fisher statistic [χ² =−2(ln p₁ + ln p₂)] to calculate a p-value for each gene evaluated against a chi-squared distribution with four degrees of freedom. Genes expressed in a concordant direction (same sign of log₂ fold change in both conditions) were assigned a signed Z-score derived from the combined p-value. Genes with discordant effects were assigned Z=0. GSEA was performed in R using the package fGSEA v.1.34.2 using Z-scores for gene ranking. Enrichment was tested against the MSigDB Reactome collection (v2025.1), significance of enrichment was determined at padj <0.05, and the top 10 most enriched Reactome pathways by absolute normalized enrichment score were plotted using ggplot2 v4.0.1.

Additional supporting raw data for this work, including all Western blot raw data and quantifications, quantification of ICC data using Imaris, and tables of DEGs from RNAseq analysis are available on Mendeley Data, V1, doi: 10.17632/fgswr9ctcr.1Mendeley Data DOI:10.17632/fgswr9ctcr.1. Park, Anna; Fish, Sarah; Samstag, Colby; Kim, Minsuh; Weiss, Jeremy; Yu, Selina; Callier, Malia; Chiu, Ella; Weiss, Joshua; Hampe, Christiane; Estes, Raja; Lin, Bernice; Khera, Arnav; Yearout, Dora; Pallanck, Leo; Zabetian, Cyrus; Davis, Marie (2026), “Dataset for’GBA1 deficiency differentially affects endolysosomal trafficking in neurons versus astrocytes.’”

## Supplemental Figure Legends

**Supplemental Figure 1**: **DNA sequencing of iPSC clones A)** DNA sequencing of *GBA1* in *GBA1^IVS/+^* iPSC clone #1 prior to CRISPR/Cas9 genome editing (top), DNA sequencing of *GBA1* in *GBA1^IVS/+^*iPSC clone A4, which underwent CRISPR/Cas9 genome editing to reverse the c.115+1 G>A, also known as IVS2+1 G>A variant back to c.115+1 G but remained unedited (middle), and DNA sequencing of *GBAP1* in *GBA1* in *GBA1^IVS/+^* iPSC clone A4 confirming no off-target genome editing of *GBAP1* (bottom). **B)** DNA sequencing of *GBA1* in *GBA1^+/+^* Rev iPSCs confirming CRISPR/Cas9 genome editing reverse the c.115+1 G>A back to G (top) and DNA sequencing of *GBAP1* in *GBA1* in *GBA1^+/+^*Rev iPSC clone confirming no off-target genome editing of *GBAP1* (bottom). **C)** DNA sequencing of *GBA1* in *GBA1^IVS/IVS^* iPSCs confirming CRISPR/Cas9 genome editing resulting in homozygous c.115+1 G>A as well as downstream intronic insertion of a T (top) and DNA sequencing of *GBAP1* in *GBA1* in *GBA1^IVS/IVS^* iPSC clone confirming no off-target genome editing of *GBAP1* (bottom).

**Supplemental Figure 2**: Confirmation of absence of chromosomal abnormalities in iPSC clones

**A)** Karyotype of *GBA1^+/+^* clone #1 iPSCs

**B)** Karyotype of *GBA1^IVS/+^* clone #1 iPSCs

**C)** Karyotype of *GBA1^IVS/+^* clone #2 iPSCs

**D)** Karyotype of *GBA1^IVS/+^* clone #6 iPSCs

**E)** Karyotype of *GBA1^+/+^* Rev clone iPSCs

**F)** Whole genome sequencing of *GBA1^IVS/+^* A4 clone iPSCs

**G)** Whole genome sequencing of *GBA1^IVS/IVS^* clone iPSCs

**H)** Whole genome sequencing of *GBA1^IVS/+^* clone #2 iPSCs

**I)** H) Whole genome sequencing of *GBA1^IVS/+^* clone #3 iPSCs

**Supplemental Figure 3: Transcriptomic profiles of *GBA1^+/+^* Rev neurons compared to *GBA1^+/+^* neurons.**

**A)** Volcano plot of global transcriptional change across *GBA1^+/+^*Rev and *GBA1^+/+^* #1,2,3 neurons. Each data point in the scatter plot represents a gene. The log2 fold change of each gene is represented on the x-axis and the log10 of its adjusted p-value is on the y-axis. Genes with an adjusted p-value less than 0.05 and a log2 fold change greater than 1 are indicated by red dots. These represent up-regulated genes. Genes with an adjusted p-value less than 0.05 and a log2 fold change less than-1 are indicated by green dots and represent down-regulated genes.

**B)** The first two principal components of principal component analysis of *GBA1^+/+^* Rev (blue) and *GBA1^+/+^* #1,2,3 neurons (red), where the x-axis is the direction that explains the most variance and the y-axis is the second most. The percentage of the total variance per direction is shown in the axis label.

**Supplemental Figure 4: Transcriptomic profiles of *GBA1^+/+^* Rev astrocytes compared to *GBA1^+/+^* astrocytes.**

**A)** Volcano plot of global transcriptional change across *GBA1^+/+^* Rev astrocytes and *GBA1^+/+^* #1,2,3 astrocytes, where each gene is plotted based on log2 fold change on the x-axis and the log10 of its adjusted p-value on the y-axis. Upregulated genes with an adjusted p-value less than 0.05 and a log2 fold change greater than 1 are indicated by red dots, and downregulated genes with an adjusted p-value less than 0.05 and a log2 fold change less than-1 are indicated by green dots

**B)** The first two principal components of principal component analysis of *GBA1^+/+^*Rev astrocytes (blue) compared to *GBA1^+/+^* #1,2,3 astrocytes (red), where the x-axis is the direction that explains the most variance and the y-axis is the second most. The percentage of the total variance per direction is shown in the axis label.

**Supplemental Figure 5: Transcriptomic profiles of *GBA1****^IVS**/+**^* **neurons or astrocytes compared to *GBA1****^IVS**/+**^* **A4 neurons or astrocytes.**

**A)** Volcano plot and PCA of *GBA1^IVS/+^*#1,2,6 neurons compared to *GBA1^IVS/+^* A4 neurons. Volcano plot of global transcriptional change across *GBA1^IVS/+^* #1,2,6 neurons compared to *GBA1^IVS/+^* A4 neurons, where each gene is plotted based on log2 fold change on the x-axis and the log10 of its adjusted p-value on the y-axis. Upregulated genes with an adjusted p-value less than 0.05 and a log2 fold change greater than 1 are indicated by red dots, and downregulated genes with an adjusted p-value less than 0.05 and a log2 fold change less than-1 are indicated by green dots. The first two principal components of principal component analysis of *GBA1^IVS/+^* #1,2,6 neurons (red) compared to *GBA1^IVS/+^* A4 neurons (blue), where the x-axis is the direction that explains the most variance and the y-axis is the second most. The percentage of the total variance per direction is shown in the axis label.

**B)** Volcano plot and PCA of *GBA1^IVS/+^*#1,2,6 astrocytes compared to *GBA1^IVS/+^* A4 astrocytes. Volcano plot of global transcriptional change across *GBA1^IVS/+^* #1,2,6 astrocytes compared to *GBA1^IVS/+^* A4 astrocytes, where each gene is plotted based on log2 fold change on the x-axis and the log10 of its adjusted p-value on the y-axis. Upregulated genes with an adjusted p-value less than 0.05 and a log2 fold change greater than 1 are indicated by red dots, and downregulated genes with an adjusted p-value less than 0.05 and a log2 fold change less than - 1 are indicated by green dots. The first two principal components of principal component analysis of *GBA1^IVS/+^* #1,2,6 astrocytes (red) compared to *GBA1^IVS/+^* A4 astrocytes (blue), where the x-axis is the direction that explains the most variance and the y-axis is the second most. The percentage of the total variance per direction is shown in the axis label.

## ACKNOWLEDGEMENTS

We thank Chris Cavanaugh, Jennifer Hesson and Julie Mathieu at the University of Washington Institute for Stem Cell and Regenerative Medicine Stem Cell Core for technical support for generating iPSCs and CRISPR/Cas9 editing of iPSCs; Jessica E. Young and members of the Young Lab at the University of Washington for technical support and advice; Darryl Hackney (Seattle VA R&D Imaging Core), Dale W. Hailey (University of Washington Garvey Imaging Core) and Nathaniel Peters (UW Keck Microscopy Center) for microscopy support. The authors especially thank the

individuals who donated tissue to generate the cell lines, without whom this work would not be possible.

## GLOSSARY

AD: Alzheimer’s disease
CBE: conduritol B epoxide
DEG: differentially expressed gene
DLB: dementia with Lewy Bodies
EEA1: early endosome antigen 1
GBA1: *glucosidase, beta acid 1*
iPSC: induced pluripotent stem cell
LAMP1: lysosome-associated membrane protein 1
MSA: multiple system atrophy
NPC: neural progenitor cell
PD: Parkinson’s disease

## CRediT Author Statement

**Anna J. Park**: Methodology, Formal analysis, Investigation, Visualization, Writing – Review & Editing

**Sarah L. Fish**: Methodology, Formal analysis, Investigation, Visualization, Writing – Review & Editing

**Colby L. Samstag**: Methodology, Formal analysis, Software, Data Curation, Writing – Review & Editing, Visualization

**Minsuh Kim**: Methodology, Formal analysis, Investigation, Visualization, Writing – Review & Editing

**Jeremy Weiss**: Software, Methodology, Formal analysis

**Selina Yu**: Investigation, Formal analysis, Visualization, Supervision, Project administration, Writing – Review & Editing

**Malia L. Callier**: Investigation, Formal analysis, Review & Editing

**Ella H. Chiu**: Investigation, Formal analysis

**Joshua Weiss**: Software, Methodology, Formal analysis

**Christiane Hampe**: Validation, Formal analysis, Review & Editing

**Raja E. Estes**: Methodology, Investigation, Review & Editing

**Bernice Lin**: Methodology, Investigation, Review & Editing

**Arnav Khera**: Investigation

**Dora Yearout**: Resources, Validation

**Leo J. Pallanck**: Resources, Writing – Review & Editing

**Cyrus P. Zabetian**: Resources, Writing – Review & Editing, Project administration, Funding acquisition

**Marie Y. Davis**: Conceptualization, Methodology, Validation, Formal analysis, Resources, Data Curation, Writing – Original draft, Writing – Review & Editing, Visualization, Supervision, Project administration, Funding acquisition

