## Supplementary figures and images for "*GBA1* deficiency differentially affects endolysosomal trafficking in neurons versus astrocytes"

### Supplemental Figure 1

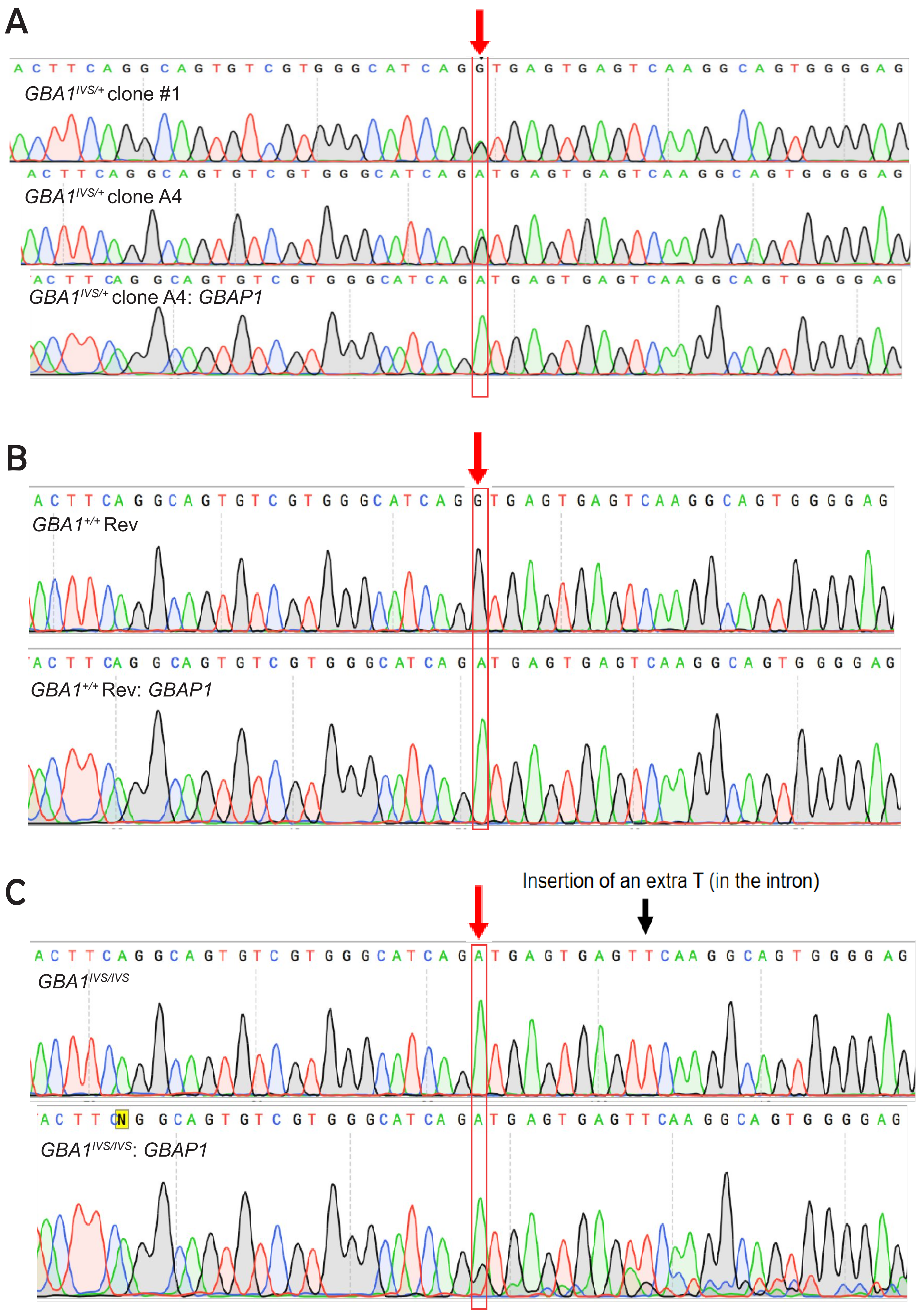

### Supplemental Figure 2

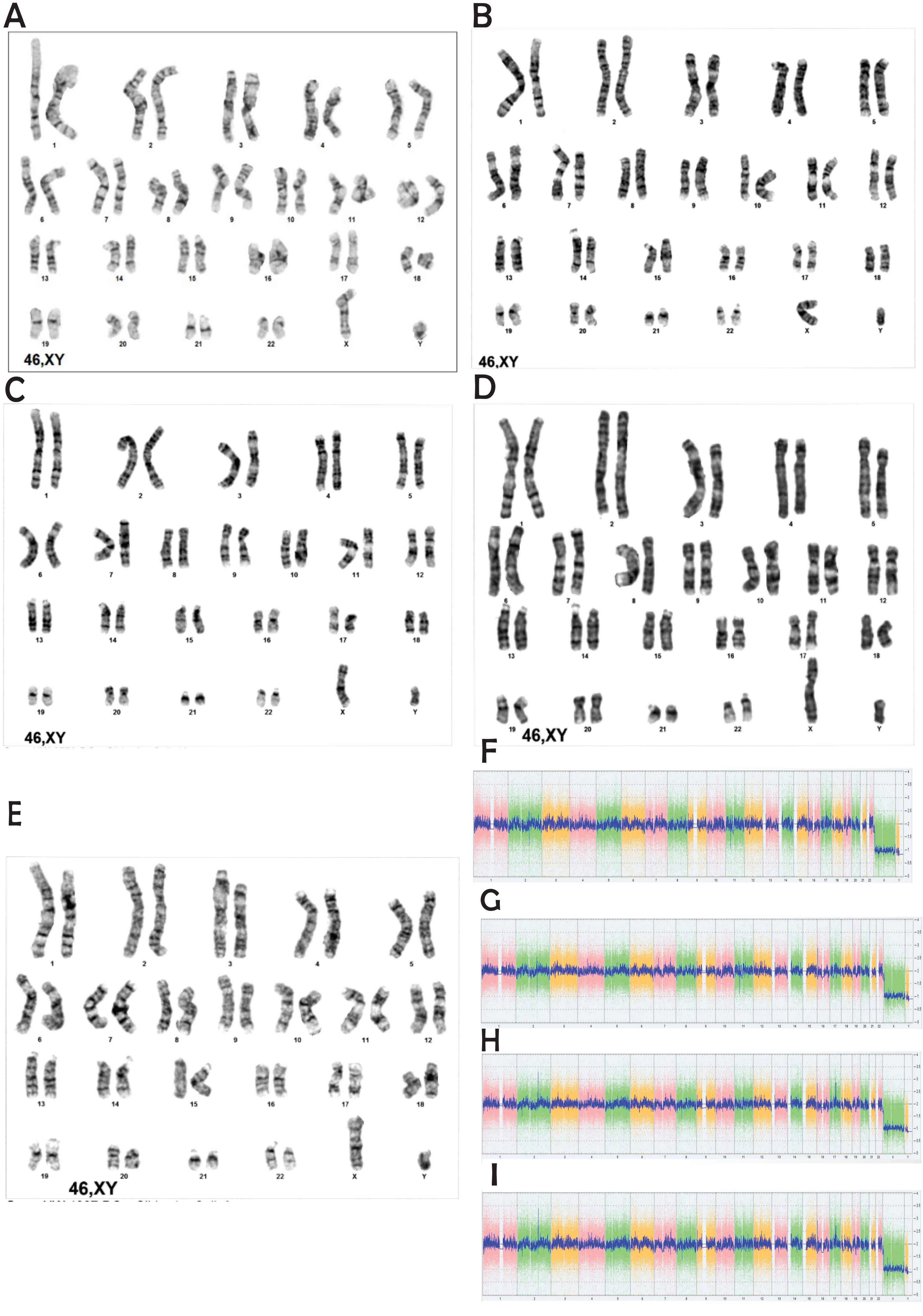

### Supplemental Figure 3

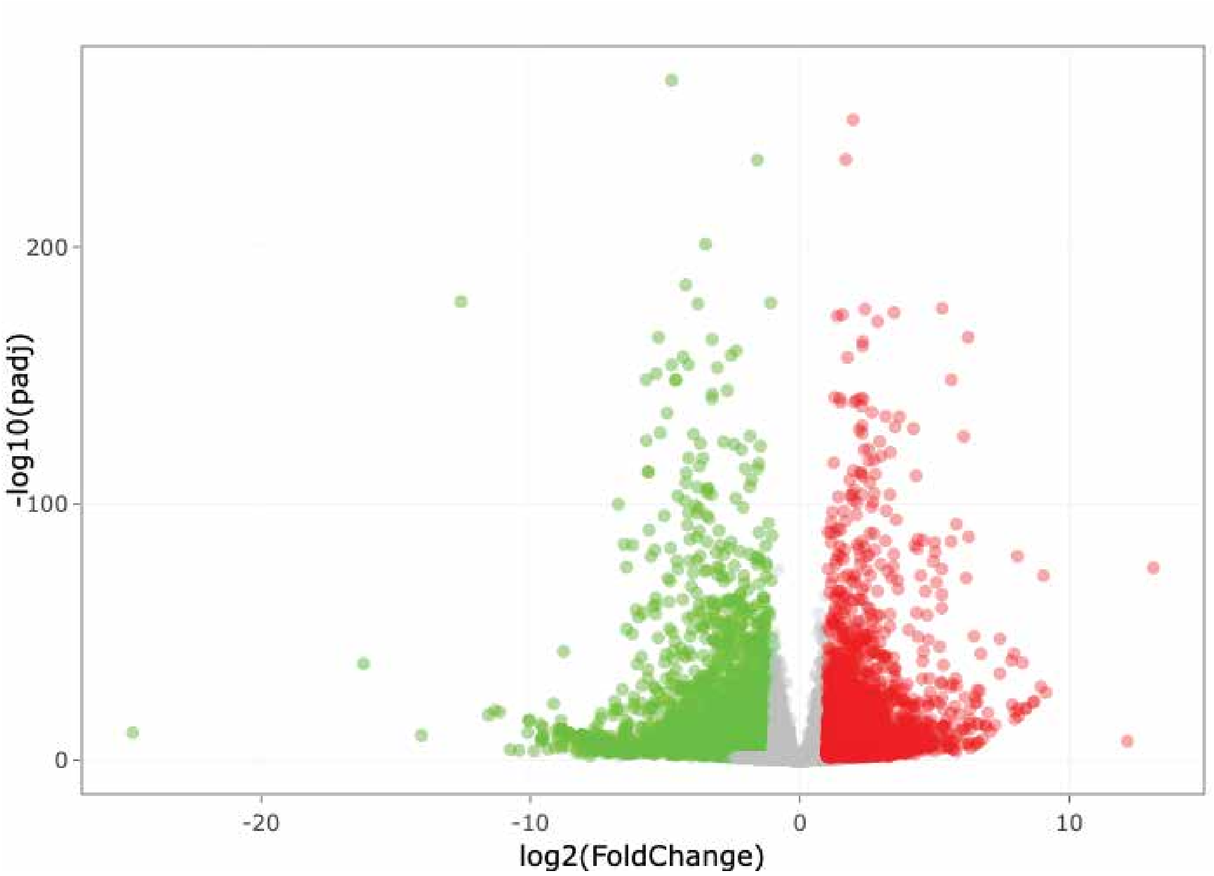

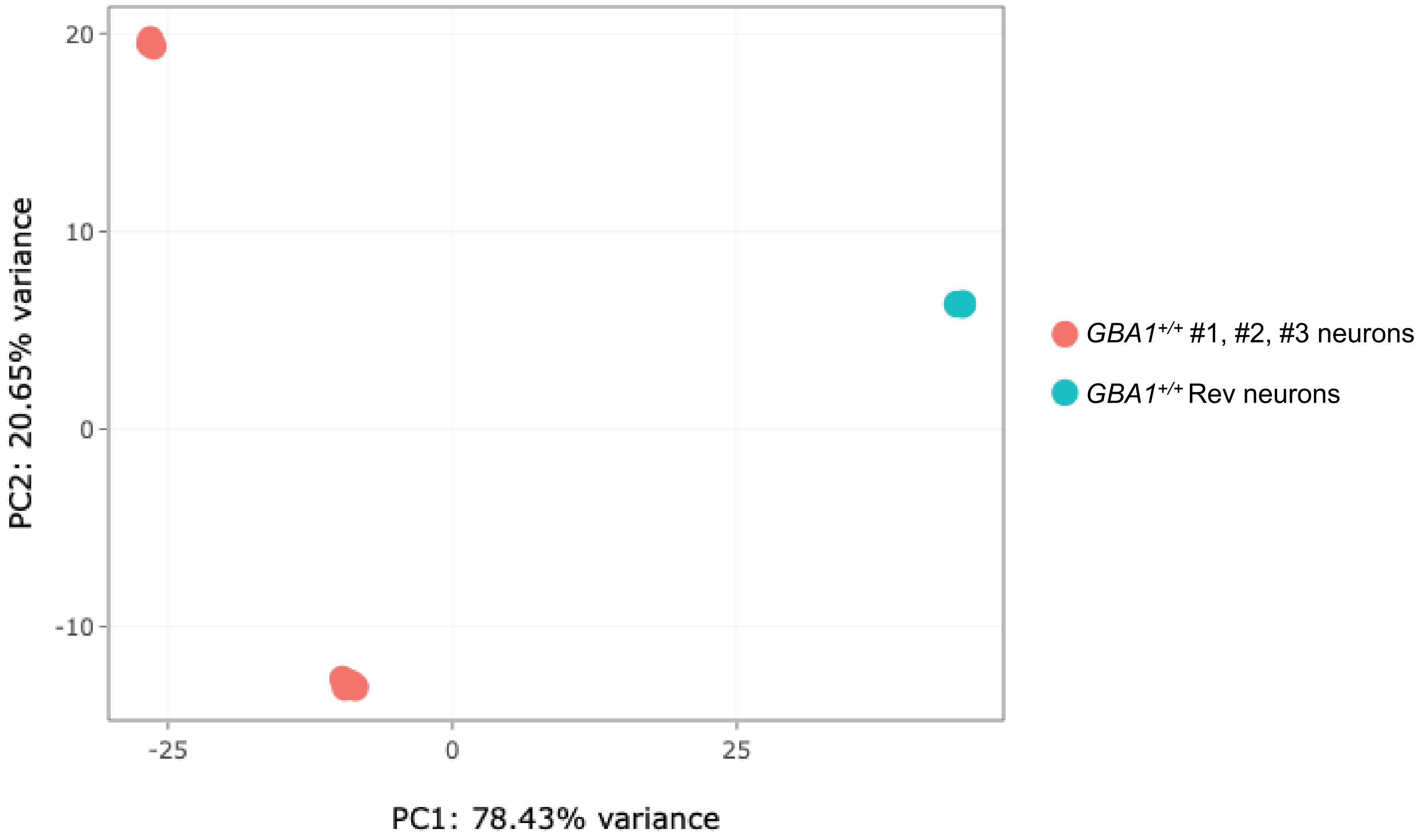

### Supplemental Figure 4

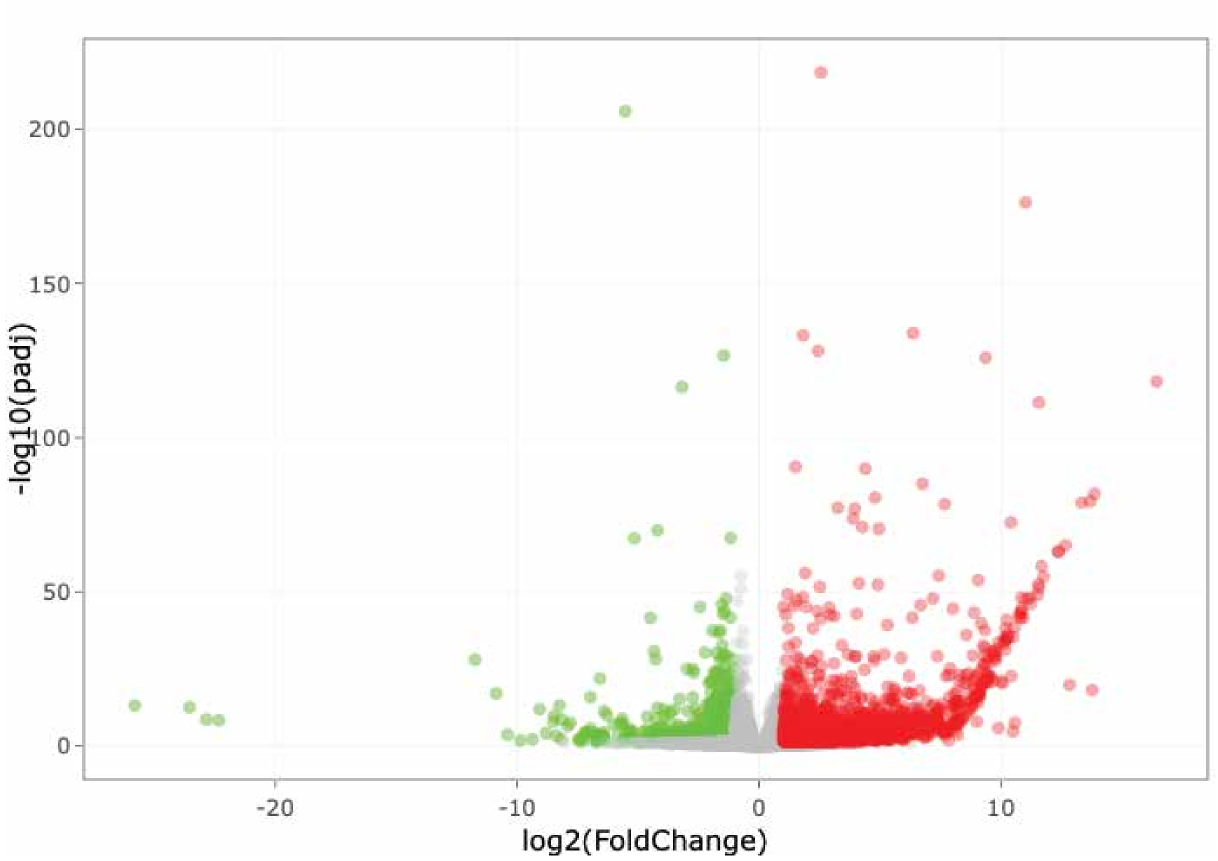

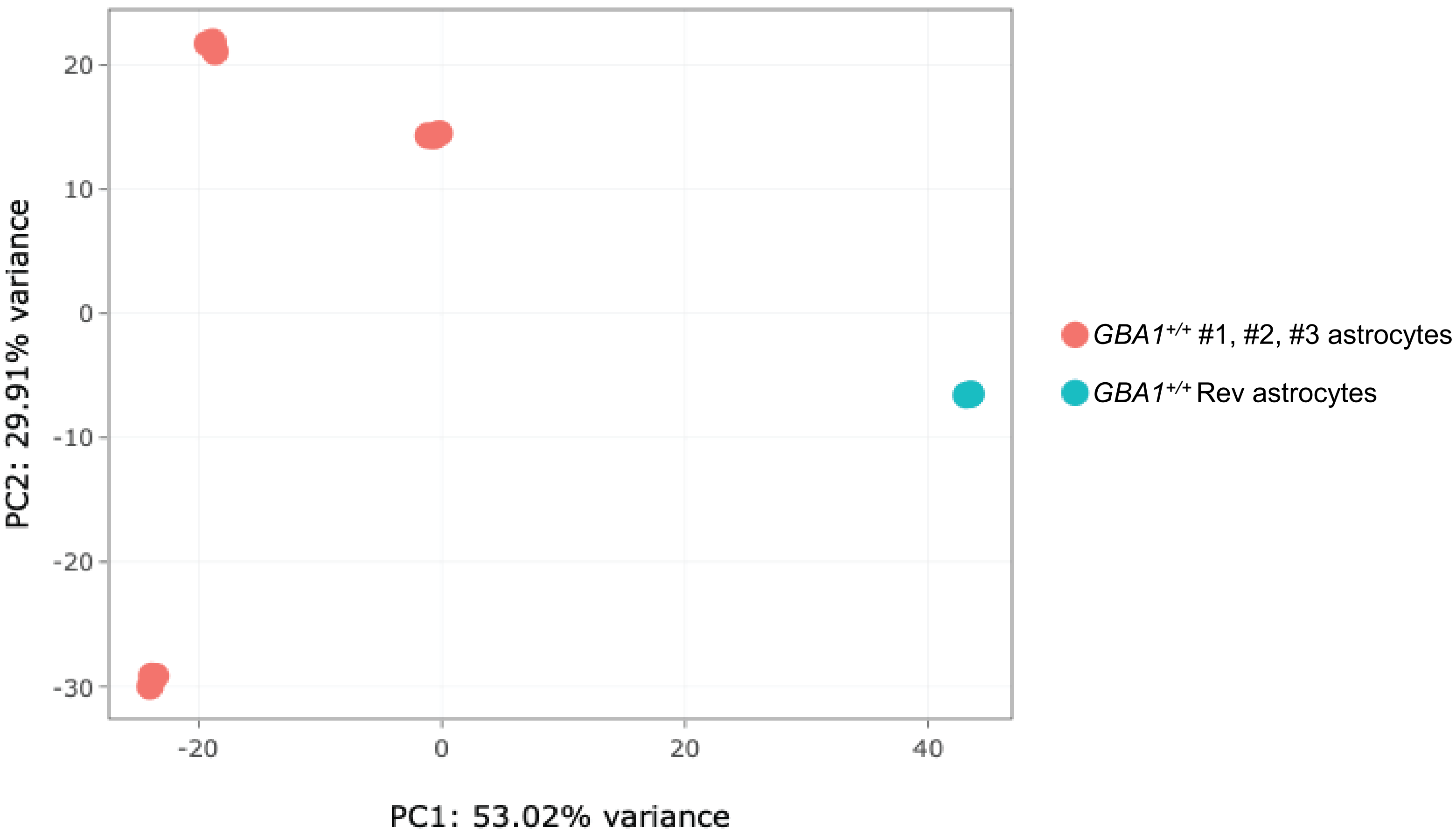

### Supplemental Figure 5

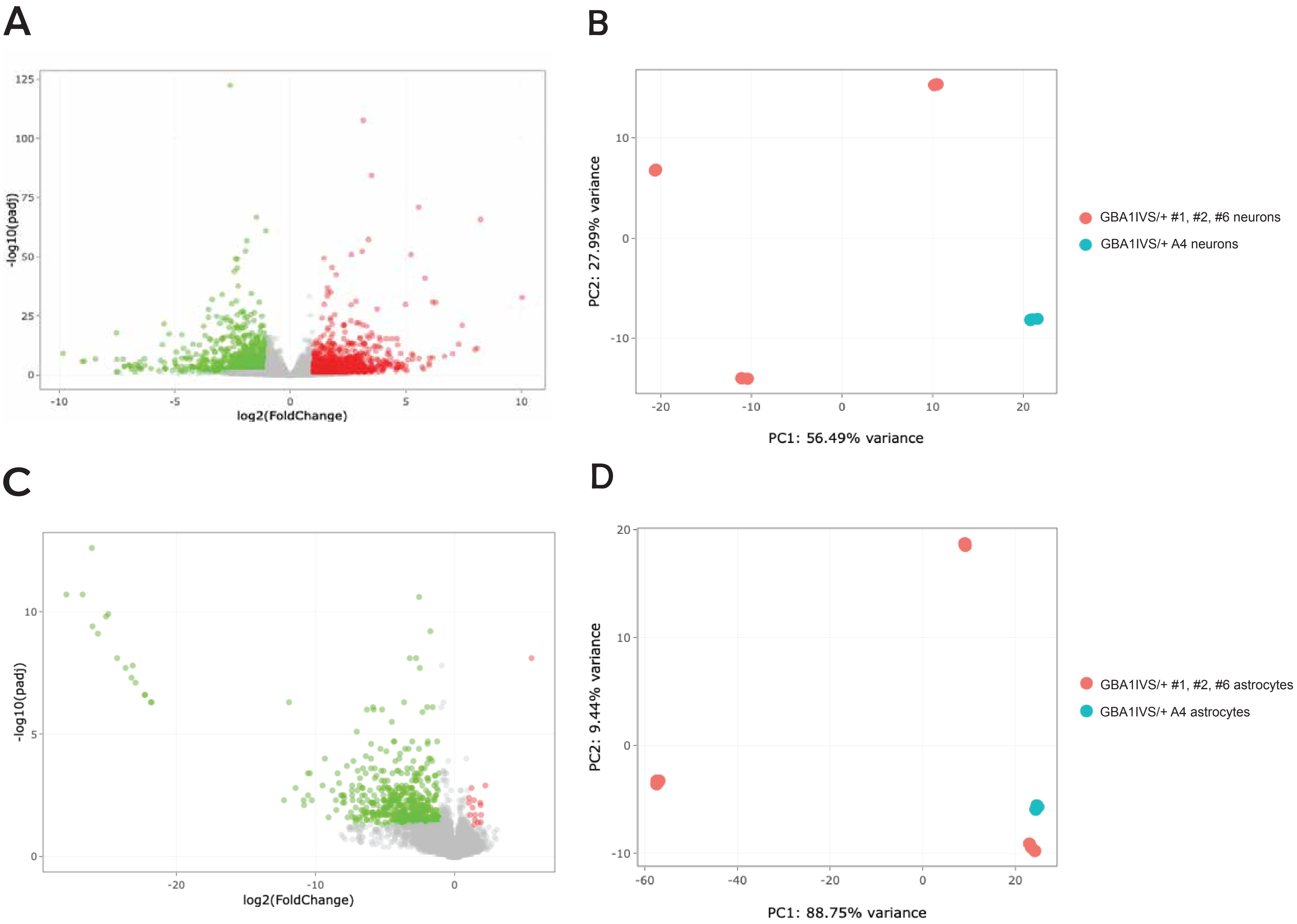
